# Transcriptional immunogenomic analysis reveals distinct immunological clusters in pediatric nervous system tumours

**DOI:** 10.1101/2022.09.20.508719

**Authors:** Arash Nabbi, Pengbo Beck, Alberto Delaidelli, Derek A. Oldridge, Sumedha Sudhaman, Kelsey Zhu, S.Y. Cindy Yang, David T. Mulder, Jeffrey P. Bruce, Joseph N. Paulson, Pichai Raman, Yuankun Zhu, Adam C. Resnick, Poul H. Sorensen, Martin Sill, Sebastian Brabetz, Sander Lambo, David Malkin, Pascal D. Johann, Marcel Kool, David T.W. Jones, Stefan M. Pfister, Natalie Jäger, Trevor J. Pugh

## Abstract

To inform immunotherapy approaches in children, we performed an immunogenomic analysis of RNA-seq data from 925 treatment-naïve pediatric nervous system tumours (pedNST) spanning 12 cancer types from three public data sets. Within pedNST, we uncovered four broad immune clusters: Pediatric Inflamed (10%), Myeloid Predominant (30%), Immune Neutral (43%) and Immune Excluded (17%). We validated these clusters using immunohistochemistry, methylation immune inference, and segmentation analysis of tissue images. We report shared biology of these immune clusters within and across cancer types, and characterization of specific immune-cell frequencies as well as T- and B-cell repertoires. We found no associations between immune infiltration levels and tumour mutational burden, although molecular cancer entities were enriched within specific immune clusters. Given the heterogeneity within pedNST, our findings suggest personalized immunogenomic profiling is needed to guide selection of immunotherapeutic strategies.

## Introduction

Cancer immunotherapies have been clinically and experimentally investigated in pediatric oncology with a wide range of response rates. Objective responses to immune checkpoint inhibitors (ICI) as a monotherapy have been limited to 5-11% of pediatric cancers (1–3). Monoclonal antibodies targeting disialoganglioside GD2 in combination with granulocyte-macrophage colony stimulating factor (GM-CSF), interleukin-2 and isotretinoin have been approved in neuroblastoma, with an overall survival rate of ∼86% (4, 5). More recently, anti-GD2 Chimeric Antigen Receptor (CAR) T-cell therapy has been investigated in diffuse midline gliomas in four patients, three of whom showed clinical improvement (6). Experimentally, novel immunotherapies have been proposed and tested in preclinical models including CAR-T targeting immune checkpoint protein, B7-H3 (7), and targeting the myeloid compartment with anti-CSF1R (8) or anti-CD47 (9). Considering the wide range of response rates for existing and emerging pediatric cancer immunotherapies, biomarkers for patient stratification are needed to identify potential candidates for clinical trials.

Immunogenomic analysis of tumours has been a major focus of biomarker discovery for immunotherapy. A prominent outcome of such studies is the FDA approval of tumour mutation burden (TMB) and microsatellite instability as the first immunotherapy-related biomarkers in adult cancers (10). Several other biomarkers have been derived from large-scale genomic and transcriptomic datasets with a major focus on adult extracranial tumours (11–15). Despite recent single-cell RNA sequencing (RNA-seq) studies in neuroblastoma (16), high-grade glioma (17), ependymoma (18, 19) and medulloblastoma (20), a comparison of immune the microenvironment across pediatric nervous system cancers and implications for informing immunotherapeutic interventions or patient selection have not been systematically analyzed. In this study, we conducted a comprehensive immunogenomic analysis of 925 treatment-naïve pediatric central and peripheral nervous system cancers to determine overall composition of the immune microenvironment and to aid in patient selection for immunotherapy clinical trials.

## Results

### Establishment of pediatric nervous system cohort for immunogenomic analysis

To study immune attributes across pediatric nervous system tumours (pedNST), we compiled a cohort of bulk RNA-seq data from three consortia: the Children’s Brain Tumour Network (CBTN), National Cancer Institute Therapeutically Applicable Research To Generate Effective Treatments initiative (NCI TARGET) (21), and the International Cancer Genome Consortium (ICGC) (22). To better represent primary pedNST, we refined the CBTN RNA-seq dataset and performed unsupervised clustering to enable comparison at the transcriptional level (n = 581, FigS1, methods). We included 195 primary samples from the ICGC (23) with matched RNA-seq, Whole Genome Sequencing (WGS) and DNA methylation data and 149 primary neuroblastomas from the NCI TARGET (21) with matched RNA-seq and Whole Exome Sequencing (WES) data (21).

The final, aggregated non-overlapping pediatric dataset for immunogenomic analysis consisted of 925 samples with primary locations in central nervous system (CNS) or peripheral nervous system : Embryonal Tumours with Multilayered Rosettes (ETMR, n = 9), Neurofibroma (NFB, n = 11), Choroid Plexus tumours (CP, n = 16), Meningioma (MNG, n = 13), Schwannoma (SCHW, n = 14), Craniopharyngioma (CPH, n = 27), Atypical Teratoid/Rhabdoid Tumours (ATRT, n = 30), Ependymoma (EPN, n = 65), pediatric High-Grade Glioma (pedHGG, n = 83), Neuroblastoma (NBL, n = 151), Medulloblastoma (MB, n = 208), and pediatric Low-Grade Glioma (pedLGG, n = 298) (Fig1A). We included data from 79 patient-derived xenograft (PDX) models as part of the Innovative Therapies for Children with Cancer Pediatric Preclinical Proof-of-Concept Platform (ITCC-P4) project as negative controls lacking immune or stromal infiltration (24, 25) (Fig1A) (TableS1).

**Figure 1.**
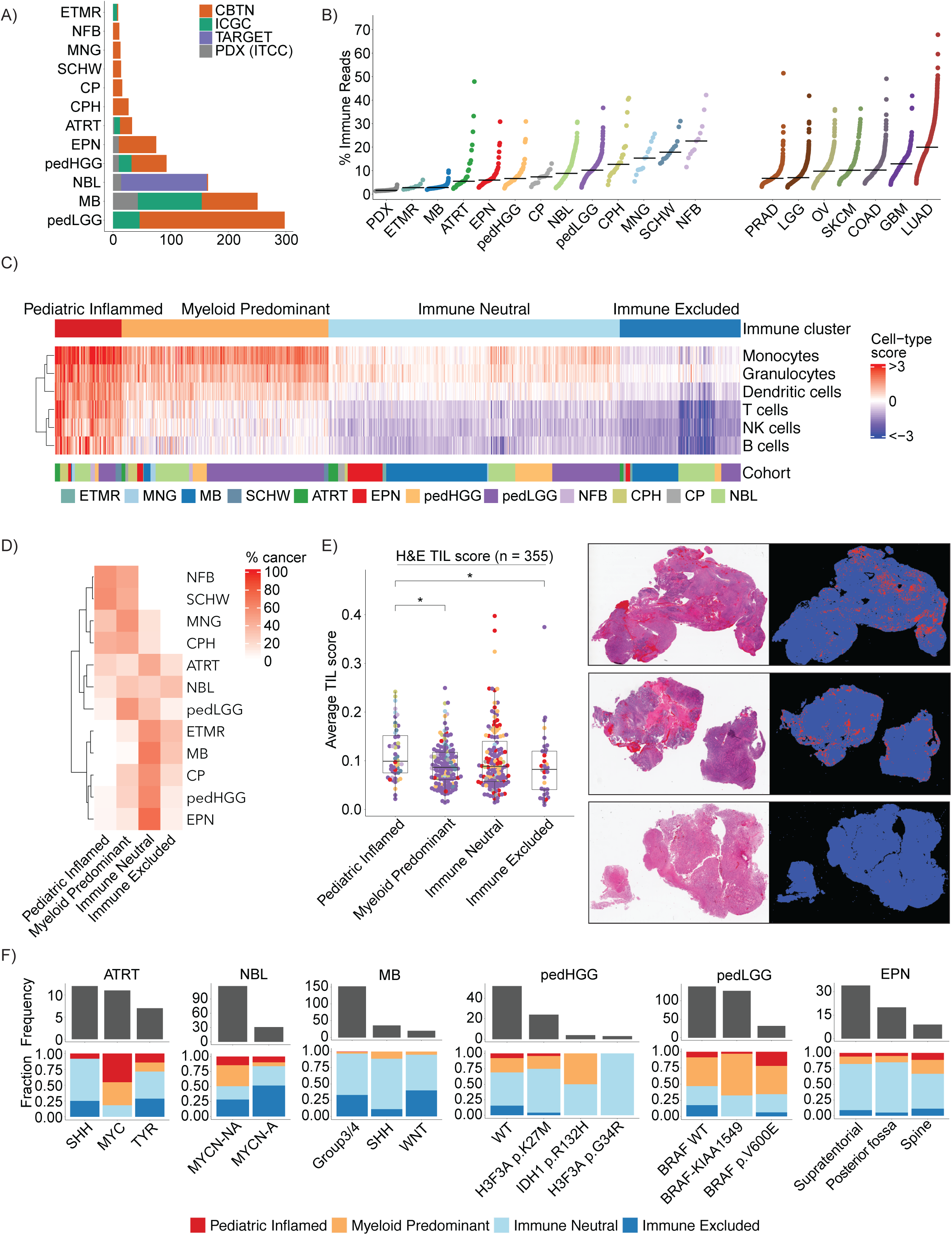
Transcriptional analysis of 925 pediatric nervous system tumours (pedNST) reveals four distinct immune clusters. **A)** Overview of cohorts and sample size in the present study (PDX: Patient-derived Xenografts, ETMR: Embryonal Tumour with Multilayered Rosettes, NFB: Neurofibroma, MNG: Meningioma, SCHW: Schwannoma, CP: Choroid Plexus tumours, CPH: Craniopharyngioma, ATRT: Atypical Teratoid/Rhabdoid Tumour, EPN: Ependymoma, pedHGG: pediatric High-Grade Glioma, NBL: Neuroblastoma, MB: Medulloblastoma, pedLGG: pediatric Low-Grade Glioma). **B)** Distribution of percentage immune reads based on ESTIMATE immune score across pediatric and adult cancers. PDX samples serve as a negative control for immune infiltration. (PRAD: Prostate Adenocarcinoma, LGG: Low-Grade Glioma, OV: Ovarian Serous Cystadenocarcinoma, SKCM: Skin Cutaneous Melanoma, COAD: Colorectal Adenocarcinoma, GBM: Glioblastoma Multiforme, LUAD: Lung Adenocarcinoma). **C)** Heatmap representing consensus clustering using enrichment scores derived from six immune-cell specific genesets across the pedNST cohort. Cell-type scores correspond to normalized gene set enrichment scores. **D)** Heatmap illustrating proportion of samples in each immune cluster across cancer entities. **E)** Boxplot showing average Tumour-Infiltrating Lymphocyte (TIL) scores as determined by segmentation analysis of pathological images across the pediatric CNS tumour samples (CBTN). Three sample images are shown representing 1% (bottom), 10% (middle) and 15% (top) TIL scores corresponding to lower quartile, mean and higher quartile, respectively. Boxes show median and interquartile range (IQR) and whiskers represent 1.5 times IQR. Two-sided rank sum test, *p < 0.05. **F)** Barplots showing frequency (top barplot) and fraction (stacked barplot) of tumour subtypes across immune clusters (SHH: Sonic Hedgehog, TYR: Tyrosine, *MYCN*-NA: *MYCN* non-amplified, *MYCN*-A: *MYCN* amplified, WT: wildtype).

### Immune infiltration analysis reveals high variability across and within cancer types

To determine the overall levels of immune infiltration across pedNST, we first assessed the performance of the deconvolution tool ESTIMATE (26) using *in silico* simulations and converted immune scores to “immune read percentage” using a non-linear regression model (FigS2). To enable comparison with adult brain tumours and cancers in which ICI agents have been clinically studied (27–31), we performed a parallel analysis of 2,452 primary adult tumour samples from The Cancer Genome Atlas (TCGA) (32, 33): Glioblastoma Multiforme (GBM, n = 153), Low-Grade Glioma (LGG, n = 507), Skin Cutaneous Melanoma (SKCM, n = 102), Colorectal adenocarcinoma (COAD, n = 298), Ovarian serous adenocarcinoma (OV, n = 373), Prostate adenocarcinoma (PRAD, n = 497), Lung adenocarcinoma (LUAD, n = 522).

Immune read percentages varied considerably within and across pedNST (median 7.2%, range 1.9%-47.9%) and in similar range compared to adult CNS tumours (median 7.8%, range 2.9%-41.8%). Common non-CNS adult cancers showed higher overall levels of immune infiltration (median 10.3%, range 2.5%-67.8%). In pedNST, highest values were in extracranial entities including neurofibroma, schwannoma and meningioma (medians 22.6%, 17.8% and 15.3%, respectively). Pediatric brain tumour entities ETMR and MB had the lowest median immune infiltration of all cancers analyzed (medians 2.7% and 2.8%) (34–36) (Fig1B). CP had consistently low variation of immune read percentage (range 3.1%-13%, median 7.3%). In contrast, ATRT exhibited the widest distribution of immune infiltration (range 2.3%-47.9%, median 5.5%), consistent with known associations between infiltration and ATRT subgroups (37, 38). Similarly, a wide range of immune read percentages were observed in CPH and NBL (range 3.9%-40.9%, median 12.6% and 2.6-30.7%, median 8.8%). These data revealed notable examples of highly-infiltrated and immune-excluded samples in pediatric nervous system tumours and identified general trends of immune infiltration.

To investigate whether the wide variability in immune infiltration could be recapitulated at the protein level, we applied immunohistochemistry (IHC) to a tissue microarray (TMA) from an independent cohort of 139 pediatric cancers, obtained from Children’s Oncology Group (COG). The median CD8 staining was greatest in ATRT followed by NBL and pedHGG, while MB and EPN had median H-score of zero (FigS3A-B). We found CD4 staining in eight samples (H-score > 0, 3 ATRTs, 3 EPN, 2 MB), while other samples showed no CD4 staining. CD19 staining was variable among cancer types ranging from 3% (MB) to 51% (NBL) (FigS3A-B). Summing H-scores across all three markers, we found highest staining in ATRT followed by NBL (medians 17 and 15) and MB showing the lowest staining. These results confirm the high variability of immune cell-type infiltration within and across pedNST, as well as generalized trends inferred from gene expression.

### Consensus gene set clustering reveals four distinct immune clusters in pedNST

In light of the variability of immune infiltrates within each tumour type, we sought to categorize immune microenvironments across pedNST. We detected unexpected immune signals inferred by various immune deconvolution algorithms when applied to PDX RNA-seq data (FigS4A-G, TableS2), which should not yield a (human) immune cell signature. This indicated wide discordance and non-specific signals across existing immune deconvolution tools. To address the non-specific immune signal that may originate from tumour cells, we sought to identify immune-cell specific genes that lack expression in pediatric nervous system cancer cells. We compiled 3,041 immune-related genes by incorporating data from four sources: ESTIMATE signature (26), an immune-cell compendium (13) and the Human Protein Atlas datasets (39, 40) (methods). We excluded genes with evidence of expression in pediatric cancer cells using data from the PDX models (ITCC-P4), single-cell RNA-seq (16,18,19) and established cell lines (CBTN). This analysis identified 791 of the 3,041 immune genes not expressed in pediatric nervous system tumour cells. We performed consensus analysis using gene sets from aforementioned immune deconvolution tools along with gene expression profiling of 28 immune cell-types (41), and blood-cell specific dataset (the Human Protein Atlas) (42). Using this approach, 216 genes were assigned to a single cell-type in at least two data types. These included genes specific for T cells (n = 41), B cells (n = 32), NK cells (n = 20), monocytes (n = 18), dendritic cells (n = 28) and granulocytes (n = 71) (TableS3). Consensus clustering of normalized enrichment scores for these cell-specific genesets identified four distinct immune clusters (Fig1C-D) that we characterized further and designated Pediatric Inflamed, Myeloid Predominant, Immune Neutral, and Immune Excluded.

To place these clusters in context of prior work in adult tumours, we applied the CRI-iAtlas adult tumour microenvironment clustering method to pedNST (14). We found that a lower proportion of pediatric brain tumours were grouped in “Immunologically quiet” or “Lymphocyte depleted” clusters compared to adult counterparts (85%, 635/749 vs 98%, 654/668). In contrast, 16.5% of pediatric extracranial tumours (NBL, NFB and SCHW, 29/176) harbored cold immune microenvironment, compared to 10.5% of adult extracranial tumours (888/8458). Across pedNST, we found 19% of pedNST (n = 174) belonged to the “Inflammatory” or “IFN-γ dominant” clusters, while 7% (n = 64) were grouped in “Wound healing”. 20% of pedLGG (60/298) were grouped in the “Inflammatory” cluster, in contrast to adult LGG, 98% of which showed a cold immune microenvironment. Almost all pedHGG samples (82/83) were grouped in the “Immunologically quiet” or “Lymphocyte depleted” clusters, similar to adult GBM (151/154). Overall, we found 72% of pedNST (n = 664) were grouped in the “Immunologically quiet” or “Lymphocyte depleted” clusters indicating a generally cold immune microenvironment in pedNST (FigS5A).

The Pediatric Inflamed cluster (n = 90, 9.7%) had the highest immune read percentage across pedNST. 57.1% of SCHW (n = 8) and 54.5% of NFB (n = 6) were clustered in this group followed by 40.7% of CPH (n = 11) and 30.8% of MNG (n = 4) (Fig1D). Five of six cases grouped in the “TGF-β dominant” cluster belonged in this cluster, followed by 42.3% of “IFN-γ dominant” samples (n = 11) (FigS5A). This cluster was devoid of samples in the “Immunologically quiet” or “Lymphocyte depleted” cluster indicating that ∼10% of pedNST samples harbor T-, B- and NK cells, monocytes, granulocytes and dendritic cells.

The Myeloid Predominant cluster (n = 279, 30.1% of pedNST) scored highly for monocyte, dendritic cell and granulocyte gene sets while harboring lower levels of lymphoid cell-types. 53.8% of MNG (n = 7) and 53% of pedLGG (n = 158) cases clustered in Myeloid Predominant followed by 44.5% of CPH (n = 12) (Fig1D). 50% of the samples within the “IFN-γ dominant” (n = 13) and 40% of the “Inflammatory” samples (n = 59) were clustered in this group (FigS5A), suggesting the inflammatory component of this cluster is driven primarily by myeloid cells.

The Immune Neutral cluster (n = 393, 42.5%) had myeloid and lymphoid cell infiltration scores near the median of the entire pedNST cohort. This cluster contained a large fraction of the cancer types with low immune read percentage including ETMR (n = 5, 55.6%), MB (n = 136, 65.4%) and EPN (n = 47, 72.3%) (Fig1D). This cluster included 56.4% of the “Immunologically quiet” (n = 62) and 48.7% of the “Lymphocyte depleted” (n = 270) tumours (FigS5A), indicating that intermediate immune infiltration across pedNST corresponds to cold immune microenvironment compared to adult cancers, which may contribute to intrinsic resistance to immune checkpoint inhibitors.

The Immune Excluded cluster (n = 163, 17.6%) received low immune inference scores for all immune cell types (Fig1C). This cluster included 32.5% of NBL (n = 49) and 29.8% of MB (n = 62) and none of the immune infiltrated cancer entities (Fig1D). Purity scores inferred from copy number alterations (43) in 156 ICGC samples revealed that samples in this cluster were of highest cancer cell content compared to Immune Neutral or Myeloid Predominant (FigS5B-C).

To validate lymphocyte infiltration levels across the immune clusters, we performed tissue image analysis for 355 pedCNS samples from CBTN with hematoxylin and eosin (H&E) stained tissue slides and matched RNA-seq (44), as well as 195 ICGC samples with matched DNA methylation arrays and RNA-seq. Consistent with the RNA-seq immune inference analysis, CP received the lowest Tumour-Infiltrating Lymphocyte (TIL) scores determined by H&E tissue analysis (median 0.08), while MNG and NFB scored highest across cancer entities (medians 0.14 and 0.11, FigS6A). TIL scores were variable across CNS tumour sites ranging from a median of 0.07 in cerebral hemispheres to a median of 0.18 in case of meninges (FigS6B). Pediatric Inflamed showed significantly higher TIL score compared to Myeloid Predominant and Immune Excluded (medians, 0.1, 0.09 and 0.08, respectively, two-sided rank sum test, p = 0.02 and 0.03, Fig1E, TableS4). The effect size was more pronounced in samples with immune read percentage > 5% (medians 0.1 and 0.03 in Pediatric Inflamed and Immune Excluded) or in those collected from cerebral hemispheres (medians 0.09 and 0.05) suggesting that high immune infiltration and composition of tumour microenvironment may influence the H&E-based inference of TILs. These findings were recapitulated by immune inference analysis utilizing the ICGC DNA methylation array data. Pediatric Inflamed tumours displayed higher levels of T-cell infiltration (pairwise two-sided Student’s t-test with Bonferroni correction, p < 0.01, FigS6D) (45). Although B-cell estimates were not significantly different between Pediatric Inflamed and Myeloid Predominant, lower levels of B cells were estimated in Immune Neutral and Immune Excluded compared to Myeloid Predominant (p < 0.001, FigS6D). Similarly, Immune Neutral and Immune Excluded showed depletion of DNA methylation-based estimates for NK cells and monocytes compared to Myeloid Predominant, in agreement with our RNA-seq clusters (p < 0.001, FigS6D).

As pedNSTs consist of distinct molecular entities and subgroups (46, 47), we next investigated associations between cancer subtypes and immune clusters (Fig1F). MYC-like ATRTs were significantly enriched in Pediatric Inflamed or Myeloid Predominant compared to non-MYC-like ATRT (one-sided Fisher’s exact test, p = 0.04 and 0.05), and 8 out of 12 SHH-like ATRT tumours clustered into Immune Neutral (one-sided Fisher’s exact test, p = 0.04) (37). In neuroblastoma (n = 148), *MYCN* non-amplified NBL cases were enriched in Myeloid Predominant (n = 41, p = 0.0008), while NBL *MYCN* amplified grouped in Immune Excluded (n = 16, p = 0.01) (48, 49). SHH MB showed a trend toward Myeloid Predominant and Immune Neutral (n = 4 and 28, p = 0.05 and 0.06), while 95% of WNT MB were grouped into Immune Neutral and Immune Excluded (19 out of 20). In pedLGG (n = 298), *BRAF* wild-type samples were enriched in Immune Excluded (n = 24, p = 4.1 x 10^-7^), *BRAF-KIAA1549* fusion-positive tumours were grouped into Myeloid Predominant (n = 82, p = 0.0004), and *BRAF* p.V600E samples were enriched in Pediatric Inflamed (n = 7, p = 0.006). In summary, our findings reveal associations between tumour-intrinsic characteristics and the immune microenvironment, although it remains to be established whether the microenvironment is sculpted by the cancer cells or whether specific cancer subtypes thrive in specific microenvironments.

We next investigated whether there were associations between patient characteristics or outcome with the immune clusters. We found an association between male sex and pedNST, more prominently seen in MB and EPN (FigS7A). A logistic regression model adjusting for cancer type indicated a trend towards an association between male genetic sex and Immune Excluded (Wald test, p = 0.05) (Fig2A). There were no significant associations between race or age and immune clusters independent of cancer type (Fig2B-C, FigS7B-C). Kaplan-Meier analysis indicated significant differences in overall survival (OS) and progression-free survival (PFS) among the immune clusters (log-rank test, p = 0.001 and 0.02) (Fig2D-E). We observed poorer OS for Immune Neutral and Immune Excluded compared to Pediatric Inflamed in a Cox proportional hazards model adjusting for cancer type and sex (Hazard Ratio (HR) = 1.56 and 1.97, Confidence Interval (CI): 1.13-2.14 and 1.39-2.79, p = 0.006 and 0.0001) (TableS5). Myeloid Predominant cases showed a trend toward improved PFS compared to Pediatric Inflamed (HR = 0.68, CI: 0.44-1.06, p = 0.08) (TableS6). Our results suggest that independent of cancer type, infiltration of specific myeloid populations provides a survival advantage over generalized hyper-infiltration of multiple immune cell types (Pediatric Inflamed) or low infiltration (Immune Neutral and Excluded).

**Figure 2.**
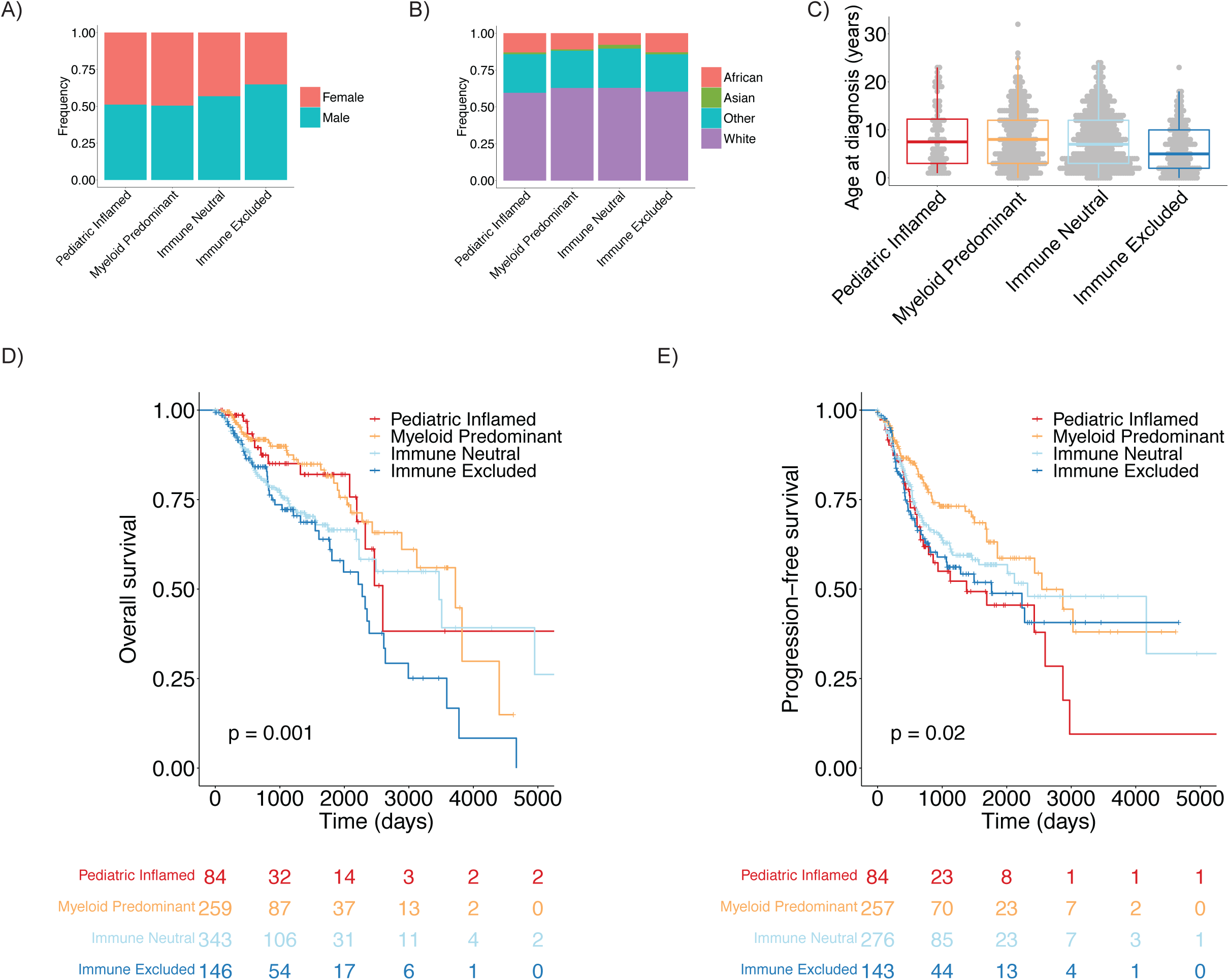
Associations of immune clusters with patients’ clinical parameters and survival. **A-B)** barplots showing distributions of gender (A) and ethnicity (B) across immune clusters in pedNST. ANCOVA, not significant. **C)** boxplot depicting age distribution across immune clusters. ANCOVA, not significant. Boxes show median and interquartile range (IQR) and whiskers represent 1.5 times IQR. **D-E)** Kaplan-Meier curves and risk tables for overall survival (D) and progression-free survival (E) among immune clusters. Log-rank test p-values are denoted.

### Gene expression analysis reveals differential molecular pathways and immunoregulatory genes in immune clusters

To understand the cellular pathways underlying each immune cluster, we conducted quantitative set analysis for gene expression (QuSAGE) (50) comparing samples in one cluster against all others while adjusting for data source and cohort. Leveraging the Molecular Signatures Database (MSigDB) Hallmark genesets (51), we found that immune-related pathways such as IFN-α, IFN-γ and allograft rejection were significantly enriched in Pediatric Inflamed and Myeloid Predominant and depleted in Immune Neutral and Immune Excluded (Fig3A, TableS7). Mitotic spindle and G2M checkpoint pathways were significantly enriched in Immune Excluded indicating a high proliferation rate with a low amounts of non-malignant cells. To investigate whether these associations could be recapitulated at the protein level, we leveraged data from 147 samples with matched RNA-seq and proteomic profiles (52). Compared to Pediatric Inflamed, Immune Neutral and Immune Excluded showed a significantly lower average protein z-score in eight immune pathways identified using QuSAGE (pairwise two-sided Student’s t-test with Bonferroni correction, p < 0.05, Fig3A-B). These findings reveal active immune pathways in Pediatric Inflamed and Myeloid Predominant consistent with the inflammatory and IFN-γ dominant adult clusters (14).

**Figure 3.**
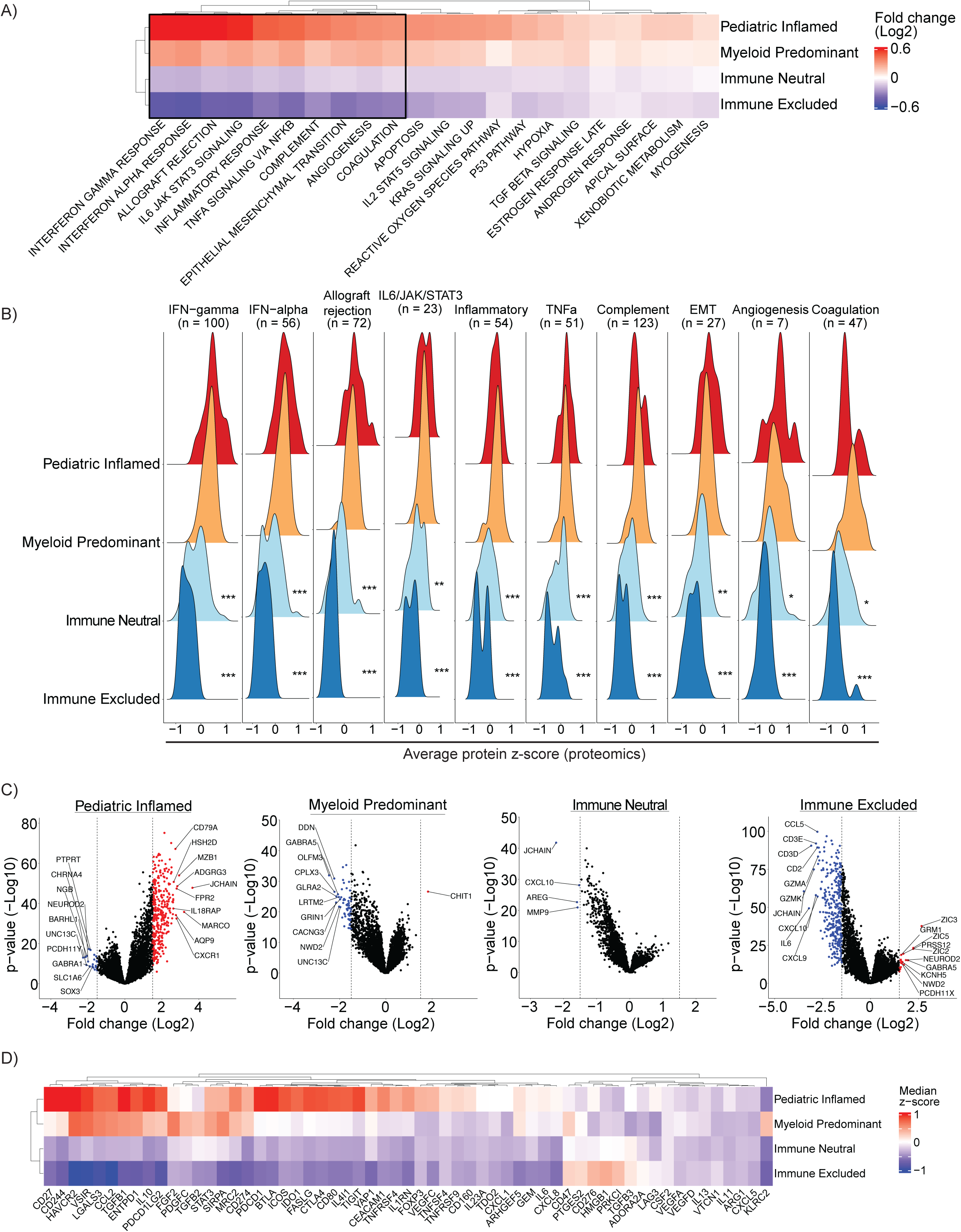
Pathway and differential gene expression analysis confirm immune features across immune clusters. **A)** Heatmap showing log-fold change in gene enrichment scores (derived from QuSAGE) in each immune cluster compared to all others in pedNST. Columns show MSigDB pathways with False Discovery Rate (FDR) < 0.1 in at least three immune clusters. Black box outlines pathways validated by protein in (B). **B)** Ridge plots illustrating average z-scores of proteins (52) involved in immune pathways from (A) across immune clusters in 141 pedCNS samples (CBTN). n indicates the number of proteins in each pathway. Pairwise two-sided Student’s t-test with Bonferroni correction, *p < 0.05, **p < 0.01 and ***p < 0.001. Significance levels are shown for comparison to Pediatric Inflamed. **C)** Volcano plots for differentially expressed genes (derived from DESeq2) in each immune cluster compared to other samples. Up-or downregulated genes with absolute log-fold change > 1.5 and FDR < 0.1 are shown in red or blue, respectively. Dashed line shows the log-fold change threshold. Top ten differentially expressed genes are annotated. **D)** Heatmap showing median z-score expression of 59 genes with known immunoregulatory functions across immune clusters in pedNST.

We further sought to determine key differential genes underlying each immune microenvironment. Using differential gene expression analysis (53), we found cell-type specific genes upregulated in Pediatric Inflamed including B-cell (*JCHAIN*, *MZB1* and *CD79A* and *MARCO*) and granulocyte specific genes (*ADGRG3* and *FPR2)* (Fig3C). In Myeloid Predominant, we found significant upregulation of *CHIT1* which encodes for chitotriosidase secreted by active macrophages (54) (Fig3C). *JCHAIN* and *CXCL10* were downregulated along with *MMP9* in Immune Neutral (Fig3C). Immune related genes such as *JCHAIN*, *CD3* chains, *IL6*, cytotoxicity genes *GZMK* and *GZMA* were significantly downregulated in Immune Excluded (Fig3C). Altogether, these results reveal core immunological pathways and genes driving immune clusters.

We sought to determine possible immunoregulatory players in pedNST. Comparison of expression levels of 59 genes with known regulatory functions (55) revealed differential expression across immune clusters after adjusting for cancer type and the total immune infiltrate (Analysis of Covariance (ANCOVA)). Members of CD28 superfamily receptor (*PDCD1*, *CTLA4*, *BTLA* and *ICOS)* were upregulated in Pediatric Inflamed (p = 6.61 x 10^-9^, 9.98 x 10^-12^, 3.04 x10^-13^ and 1.67 x 10^-17^, respectively, Fig3D). Genes encoding PD-L1 and PD-L2 (*CD274* and *PDCD1LG2*) were upregulated in Pediatric Inflamed (p = 0.03 and 6.34 x 10^-4^). Genes involved in regulatory T-cell (Tregs) function and activation (*IDO1*, *STAT3* and *FOXP3*) were upregulated in Pediatric Inflamed (p = 1.63 x 10^-8^, 0.002 and 2.31 x 10^-11^, respectively). These findings reveal high expression levels of immune checkpoint genes as well as genes involved in immunosuppression in Pediatric Inflamed.

In Myeloid Predominant, *TGFB3* encoding for TGF-β3 was significantly upregulated (ANCOVA, p = 0.007) (Fig3D). *LGALS3* encoding for Galectin-3 with negative regulatory functions in macrophages (56) was upregulated in Myeloid Predominant suggesting the prominent macrophage influence in this cluster (p = 0.004). Other myeloid-specific genes with immunoregulatory functions were upregulated in both Pediatric Inflamed and Myeloid Predominant and included *IL4I1* (57)*, CCL2* (58) and *FGF2* (59) (Fig3D). These findings show preferential expression of myeloid-related regulatory genes in Myeloid Predominant.

In Immune Excluded, we found that *CD276* encoding immune checkpoint protein B7-H3 to be highly expressed (p = 9.61 x 10^-4^, Fig3D). Unexpectedly, among immune checkpoint genes, *LAG3* was expressed at higher levels in Immune Excluded (p = 0.004). Further inspection of immune checkpoint genes in PDX data confirmed high expression of *LAG3* (FigS8A). Across MB samples in Immune Excluded, WNT subgroup expressed *LAG3* at significantly higher levels compared to SHH and there was a non-significant trend between WNT and Group3/4 (two-sided Student’s t-test, p = 0.02 and 0.08, FigS8B). These results suggest *LAG3* expressed by tumour cells in pedNST and identify B7-H3 as a possible mechanism of immune exclusion in subsets of pedNST.

In neuroblastoma, profiling expression of five immune checkpoint genes revealed ∼10% of samples had exclusively elevated expression of *HAVCR2* gene encoding TIM3 (FigS8C). We confirmed this finding using a non-overlapping NBL dataset of 209 immune infiltrated tumours. In this cohort, we found 13 cases (6.2%) that showed elevated expression of *HAVCR2* and low expression of *LAG3* (FigS8C). Immunohistochemical staining for TIM3 and LAG3 proteins using an independent TMA showed that TIM3 was detectable in 14 samples without LAG3 staining (FigS8D). These findings identify an *HAVCR2* expression in a subset of neuroblastomas suggesting a distinct mode of immunosuppression.

### Immune microenvironment associations with tumour intrinsic genomic alterations and tumour mutation burden

We next asked whether immune clusters are associated with tumour mutation burden (TMB). We did not find a statistically significant difference in the total number of non-synonymous single-nucleotide variants (SNV) per coding megabase (SNV/Mb) or SNV + Insertion/deletion (Indel) /Mb (SNV + Indel /Mb) across immune clusters (ANCOVA comparing to Pediatric Inflamed, Fig4A-B, TableS9-S10). However, when we looked at 63 pedHGG samples, we found Myeloid Predominant and Immune Neutral harbored significantly higher TMB compared to Immune Excluded (two-sided rank sum test, p = 0.02, Fig4C). Four of 6 cases with > 5 SNV + Indel/Mb and germline variations in *MLH1*, *MSH2*, *MSH6*, *PMS2*, *POLE* or *POLD1* belonged to Myeloid Predominant, suggesting higher levels of immune infiltration in patients with biallelic Mismatch Repair Deficiency (bMMRD) syndrome. Our findings otherwise revealed no associations between the immune microenvironment and TMB in non-hypermutant pedNST, suggesting TMB, with the exception of MMR-associated tumours, may not be an appropriate biomarker for immune checkpoint inhibitors in this population.

**Figure 4.**
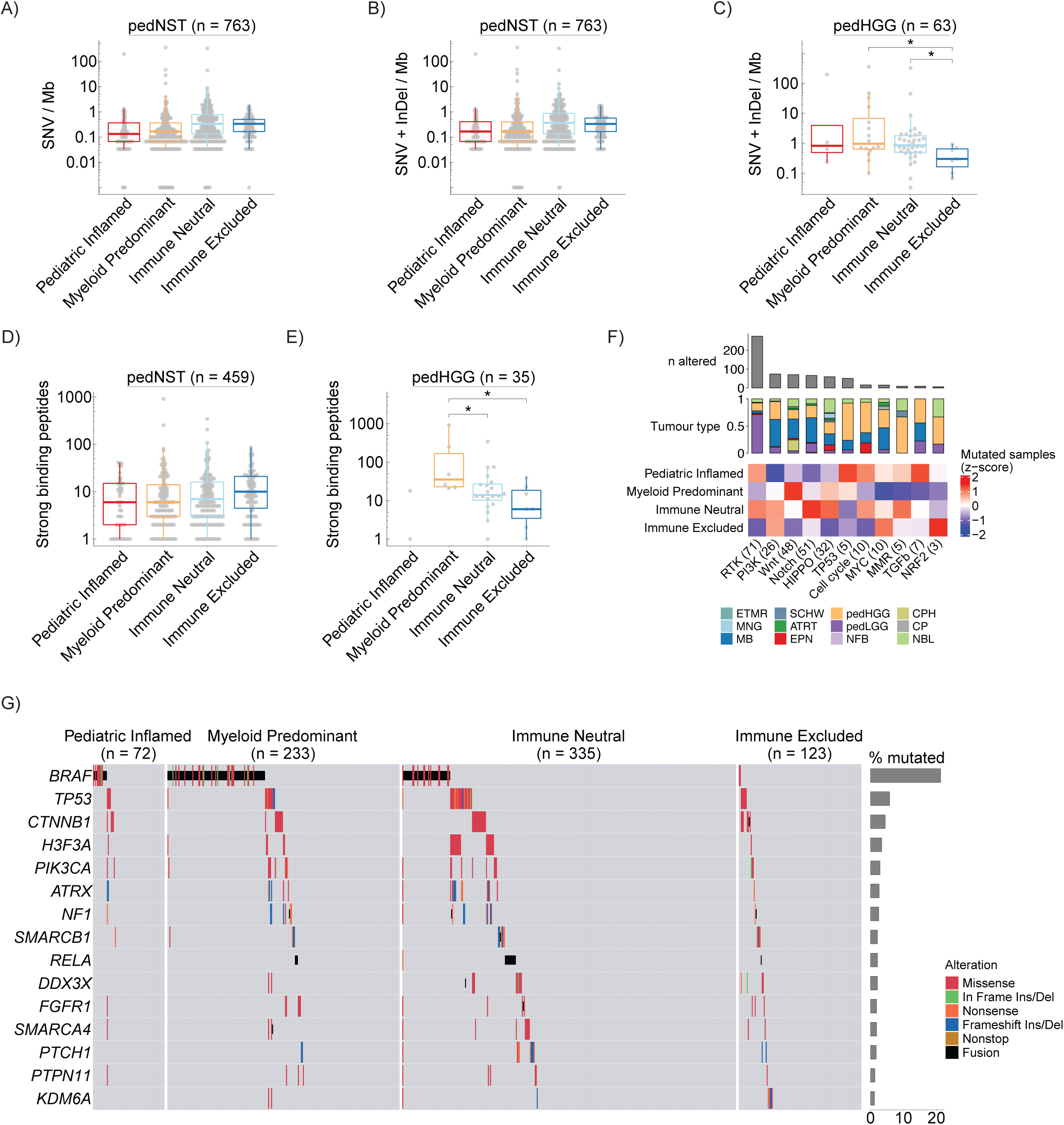
No associations between TMB or predicted neopeptides across immune clusters. **A-B)** Boxplots showing TMB defined as SNV per megabase (Mb) (A) or SNV + Indel per Mb (B) across immune clusters in pedNST. ANCOVA, not significant. **C)** Boxplot showing TMB for pedHGG samples across immune clusters. Two-sided rank sum test, *p < 0.05. **D)** Boxplot showing number of predicted strong binding peptides (defined as binding affinity ≤ 0.5) for pedNST samples across immune clusters. ANCOVA, not significant. **E)** Boxplot showing number of predicted strong binding peptides for pedHGG samples across immune clusters. Two-sided rank sum test, *p < 0.05. **F)** Heatmap illustrating scaled number of samples (z-score) with at least one non-synonymous SNV/Indels in genes involved in oncogenic pathways, as defined by TCGA. Barplot shows the total number of samples with alterations in each pathway. Stacked barplot shows proportion of tumour types present in samples with alterations in each pathway. Numbers in brackets indicate the number of altered genes in each pathway. **G)** Oncoprint illustrating the distribution of somatic mutations in the top 15 most commonly altered genes in pedNST across the four immune clusters. In all boxplots, boxes show median and IQR and whiskers represent 1.5 times IQR.

To investigate the MHC presentation potential of somatic mutations in pedNST, we determined patients’ HLA class I types (60) (FigS9) and performed the mutant peptide extractor and informer (MuPeXI) (61) analysis. We identified 7,591 strong binding and 21,680 weak binding peptides, as defined previously (62). The number of predicted strong binding peptides were highest in NBL and pedHGG (medians 19 and 17) and lowest in CPH (median 2.5), consistent with its low TMB (median 0.07 SNV + Indel/Mb, FigS10A-B). Differences in the number of strong or weak binding peptides did not reach statistical significance after adjusting for cancer type (ANCOVA, Fig4D, FigS10C, TableS11). Consistent with our TMB observation, pedHGG showed a significantly higher number of strong or weak binding peptides in Myeloid Predominant compared to Immune Neutral or Immune Excluded (Fig4E, FigS10D). These data indicate neopeptides are not independently linked to immune infiltration in pedNST, which is consistent with the lack of association between TMB and the immune microenvironment.

We hypothesized that disruptions in oncogenic pathways may be associated with distinct immune clusters. We identified samples with alterations (SNVs, Indels and fusions) in at least one gene in the ten TCGA oncogenic pathways (63) (Fig4F). Tumours with somatic alterations in members of the Receptor Tyrosine Kinase (RTK) pathway were most frequently found in Myeloid Predominant (Cochran-Mantel-Haenszel (CMH) test, p = 6.1 x 10^-5^, Fig4F). Pediatric Inflamed showed a higher percentage of samples with mutations in the Mismatch Repair (MMR) pathway, although this difference did not reach statistical significance across cancer entities (p = 0.41, Fig4F). Among common driver mutations (Fig4G) (64), we found that *BRAF*-altered samples were differentially clustered among immune clusters, with *BRAF-KIAA1549* fusion-positive samples grouped primarily in Myeloid Predominant (p = 3 x 10^-6,^ Fig4G). While the causal relationship between tumour-intrinsic genomic alterations and immune microenvironment remains unclear, our results reveal associations of altered molecular pathways and microenvironmental features.

### T- and B-cell repertoire analysis suggest antigen presentation and clonal outgrowth in immune infiltrated pedNST

We sought to determine the extent of clonal diversity for T and B cells across pedNST. Using an immune repertoire processing framework (65), we recovered a total of 23,842 Complementarity-Determining Regions (CDR3β) sequences from 582 pedNST samples. To validate the diversity estimates, we used the CapTCR-seq method (66) to enrich T-Cell Receptor (TCR) sequences in RNA-seq libraries from adult and pediatric cancer samples (n = 26). TCRβ estimated Shannon diversity inferred from bulk RNA-seq showed the highest correlation with the observed Shannon diversity obtained from CapTCR-seq (Fig5A, FigS11A-B). Comparing the diversity estimates and total number of TCRβ reads showed a linear correlation (adjusted r^2^ = 0.72, Fig5B). However, 12 samples displayed lower diversity relative to their number of TCRβ reads, suggesting a T-cell clonal expansion in these samples. Conversely, eight samples harbored outlier high diversity indicating several individual clones infiltrating these tumours (Fig5B). These findings indicate a reliable method of T-cell diversity inference and identification of potential polyclonal and clonal repertoires using bulk RNA-seq data.

**Figure 5.**
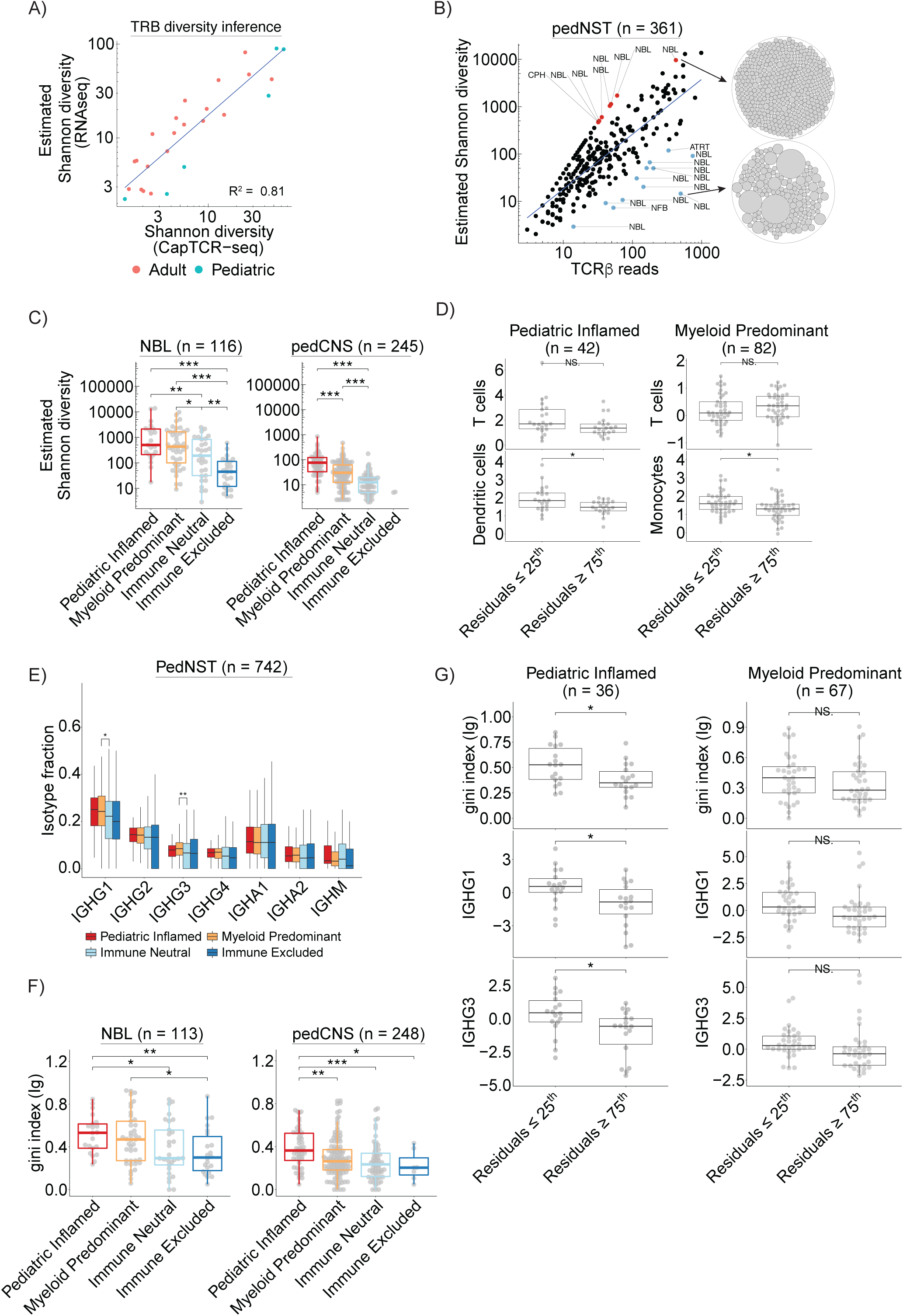
T-cell repertoire analysis reveals associations of T-cell clonal expansion with B-cell repertoire and immune microenvironment in Pediatric Inflamed. **A)** Scatterplot depicting a linear correlation between TCRβ Shannon diversity estimated from RNA-seq and measured by capturing all TCR sequences from the same RNA-seq libraries in adult and pediatric cancer samples using CapTCR-seq**. B)** Scatterplot showing correlation between TCRβ estimated Shannon diversity and total number of TCRβ reads. Blue line shows fitted linear regression. Red and blue dots represent polyclonal and clonal T-cell repertoires defined as residuals greater than two absolute standard deviations. Circle plots to the right illustrate two examples of polyclonal (top) and clonal (bottom) T-cell repertoires. Each circle is one T-cell clone and circle diameters are proportional to TCRβ reads. **C)** Boxplots showing differences in log-transformed estimated TCRβ Shannon diversity across immune clusters for NBL (left) and pedCNS (right). Two-sided Student’s t-test, *p < 0.05, **p < 0.01 and ***p < 0.001. **D)** Boxplots comparing levels of T cells, dendritic cells or monocytes, as determined in Fig1C, in samples with TCRβ residuals (obtained from the linear regression in (B)) ≤ 25th percentile and ≥ 75th percentile of Pediatric Inflamed (left) or Myeloid Predominant (right). Two-sided Student’s t-test, *p < 0.05, NS: not significant. **E)** Boxplot showing proportion of specific immunoglobulin isotypes in B-cell repertoires across immune clusters in pedNST. Two-sided Student’s t-test, *p < 0.05 and **p < 0.01. **F)** Boxplots comparing immunoglobulin clonality (gini index) across immune clusters for NBL (left) and pedCNS (right). Two-sided Student’s t-test, *p < 0.05, **p < 0.01 and p < 0.001. **G)** Boxplots comparing levels of gini index or tumour-type normalized expression of *IGHG1* or *IGHG3* in samples with TCRβ residuals (obtained from the linear regression in (B)) ≤ 25th percentile and ≥ 75th percentile of Pediatric Inflamed (left) or Myeloid Predominant (right). Two-sided Student’s t-test, *p < 0.05, NS: not significant. In all boxplots, boxes show median and IQR and whiskers represent 1.5 times IQR.

Using estimated Shannon diversity, we analyzed 361 pedNST samples for TCRβ diversity (FigS11C, methods). NBL samples in Pediatric Inflamed showed significantly higher diversity compared to NBL samples in Immune Neutral and Excluded (two-sided Student’s t-test, p = 0.006 and 4.3 x 10^-7^, Fig5C). Within pedCNS, Pediatric Inflamed had higher diversity compared to Myeloid Predominant and Immune Neutral (p = 5.5 x 10^-8^ and < 2.2 x 10^-16^, Fig5C). To determine microenvironmental changes that co-vary with T-cell diversity independent of T-cell infiltration, we calculated residuals from a linear regression model for TCRβ diversity and reads (Fig5B). In Pediatric Inflamed, samples with residuals in lower quartile, corresponding to uneven T-cell repertoire, had significantly higher dendritic cell scores (p = 0.01, Fig5D). Conversely, within Myeloid Predominant, samples with residuals in lower quartile harbored significantly higher levels of monocytes (p = 0.03, Fig5D). These results reveal associations of myeloid compartment with clonal T-cell repertoire, suggesting the possibility of clonal outgrowth as a consequence of interactions with antigen presenting cells.

We used a similar immune repertoire inference tool as for the T-cell repertoire analysis (65) and recovered 197,769 unique immunoglobulin heavy chain (IGH) CDR3 sequences. Across pedNST, we found the highest number of IGH isotypes in CPH and NBL samples (FigS11D). Consistent with findings in adult cancers (67), IGHG1 constituted the largest proportion of B-cell repertoire relative to total number of isotypes (FigS11D). We found IGHG1 and IGHG3 were significantly enriched in Myeloid Predominant compared to Immune Neutral and Immune Excluded (two-sided Student’s t-test, p = 0.03 and 0.008, Fig5E), suggesting that potent antibody-producing cells infiltrated these tumours.

To determine the extent of B-cell clonal expansion in pedNST that may suggest antigen recognition (68), we used the gini index as a measure of uneven B-cell cluster distribution in each sample (69) (FigS11E, methods). Across NBL samples, Immune Neutral and Excluded had significantly lower gini index compared to Pediatric Inflamed (medians 0.29, 0.3 and 0.53, respectively, two-sided Student’s t-test, p = 0.04 and 0.009, Fig5F). Within pedCNS, Pediatric Inflamed had the highest median gini index compared to other clusters (medians 0.36, 0.26, 0.23 and 0.2 for Pediatric Inflamed, Myeloid Predominant, Immune Neutral and Excluded, respectively, p = 0.005, 0.0003 and 0.02, Fig5F). Within Pediatric Inflamed, samples with clonal T-cell repertoire (residuals ≤ 25^th^) also had clonal B-cell repertoire and expressed higher levels of *IGHG1* and *IGHG3* compared to samples with polyclonal T-cell repertoire (residuals ≥ 75^th^) (p = 0.02, 0.01 and 0.01, Fig5G). We did not find this association in Myeloid Predominant that harbored lower levels of T cells (Fig5G), suggesting that both the extent and clonality of B-cell infiltration may contribute to T-cell clonal expansion. Further characterizing lymphoid and myeloid subgroups in Pediatric Inflamed and Myeloid Predominant provided mechanistic cues to immune dysfunction and evasion (Supplementary Notes, FigS12). These results suggest the interplay between T- and B-cell compartments in pedNST and may provide mechanistic insights into antigen presentation and clonal outgrowth.

## Discussion

We report universal immune microenvironment groups across 12 cancer entities highlighting tumour-agnostic immunological similarities in pediatric primary nervous system tumours. Across our compendium, we found ∼72% of samples harbored generally cold immune microenvironments in higher frequency compared to adult tumour microenvironments (14). Within our cohort, we found 10% of samples showed an inflamed microenvironment harboring high levels of lymphoid- and myeloid-cell infiltration and diverse T- and B- cell repertoire. In this cohort of non-hypermutant cancers, TMB was not independently associated with an inflamed microenvironment. This finding is in contrast to bMMRD cancers where samples with high CD8+ T-cell infiltration had higher SNV/Mb (70). Although we could not directly study the associations between TMB and ICI response, our findings do not support TMB as a biomarker for use in non-hypermutant pediatric cancers.

Analysis of immunoregulatory genes and immune cell subtypes provided insights into the ligand-receptor interactions with translational implications. In Pediatric Inflamed, a number of upregulated genes overlapped with those differentially expressed in post-treatment samples collected from melanoma patients responding to nivolumab (71). These include immune checkpoint genes (*PDCD1, TIGIT, CTLA4* and *BTLA*) as well as genes involved in T-cell cytotoxic functions (*IL4I1, FASLG, TNFRSF9, TNFRSF4, CD244, CD27, CD80* and *ICOS*) and suggest samples in this cluster may be good candidates for ICI therapy. The Myeloid Predominant cluster showed elevated expression of two immunoregulatory genes with myeloid-specific functions: *TGFB1* (72, 73) and *SIRPA*, encoding for SIRPα that negatively regulates phagocytosis (74). Blocking this interaction with anti-CD47 promotes cell killing in preclinical models (9) and a phase I clinical study testing its efficacy is ongoing (NCT02216409). Our results suggest disrupting cellular interactions involving the lymphoid and myeloid compartments may be beneficial among a subset of pedNST to reshape the tumour microenvironment and elicit anti-tumour immunity.

In conclusion, we identified distinct immune microenvironment clusters across pediatric nervous system tumours and proposed microenvironmental mechanisms of immune dysfunction and suppression. With immunotherapy becoming more widely available in the pediatric oncology armamentarium, our findings highlight the value of immunogenomic approaches to guide patient stratification and inform precision oncology programs.

## Methods

### Human subjects

We curated a total of 925 tumours from the Children’s Brain Tumour Network (CBTN, n = 581), Therapeutically Applicable Research To Generate Effective Treatments (NCI TARGET, n = 149) and the International Cancer Genome Consortium (ICGC, n = 195). We selected tumours from pediatric patients (median age 7 years) who had RNA-seq data generated from their primary tumours. We focused this study to major types of pediatric CNS tumours and neuroblastoma; ETMR (n = 9), NFB (n = 11), CP (n = 16), MNG (n = 13), SCHW (n = 14), CPH (n = 27), ATRT (n = 31), EPN (n = 65), pedHGG (n = 83), NBL (n = 151), MB (n = 208), and pedLGG (n = 298). As adult cancer comparator, we included TCGA participants with common types of adult cancers; Glioblastoma Multiforme (GBM, n = 153), Low-Grade Glioma (LGG, n = 507), Skin Cutaneous Melanoma (SKCM, n = 102), Colorectal adenocarcinoma (COAD, n = 298), Ovarian serous adenocarcinoma (OV, n = 373), Prostate adenocarcinoma (PRAD, n = 497), and Lung adenocarcinoma (LUAD, n = 522).

### Tumour datasets

We used RNA-seq datasets from the TARGET (https://ocg.cancer.gov/programs/target), CBTN (https://cbtn.org/research/specimendata/), ICGC (https://icgc.org/icgc/cgp/62/345/822) and TCGA (https://www.cancer.gov/about-nci/organization/ccg/research/structural-genomics/tcga) in this study. For the ICGC dataset, we chose 195 primary CNS tumour samples with matched RNA-seq and WGS, and complete clinical data (23). For the TARGET dataset, we chose 149 primary neuroblastoma samples with matched RNA-seq and WES. For the CBTN dataset, we refined the original dataset (n = 996) and excluded data from 22 cell lines, 153 progression samples, 70 recurrences and 16 secondary malignancies. We further removed 98 samples from rare (defined as ≤ 10% of the cohort) or unannotated tumours. To enable comparison at the transcriptional level, we performed unsupervised clustering on primary samples and removed 51 samples that did not match their corresponding pathological annotation (FigS1A-B). This sample curation resulted in 581 samples from the CBTN consortium (FigS1C). We further annotated medulloblastoma and ATRT tumour subtypes using unsupervised clustering for downstream analysis (FigS1D-E). These resulted in a cohort of 925 pediatric CNS tumours and neuroblastomas.

We accessed CBTN, TARGET and TCGA data through the Kids First Data Resource Centre (https://portal.kidsfirstdrc.org/) and used the CAVATICA platform for data processing and analysis (https://cavatica.sbgenomics.com/). In total, we analyzed 925 cases of pediatric nervous system tumours (CNS and neural crest tumours) (CBTN, TARGET and ICGC). We included RNA-seq data from 79 PDX models (2 ATRT, 10 EPN, 10 HGG, 14 NBL, 43 MB) obtained from ITCC as control for immune infiltration and immune-cell specific geneset analysis (https://www.itcc-consortium.org/). We used an NBL dataset as a validation set for immune checkpoint profiling that was available on CAVATICA (phs001436.c1). This dataset was not included in the pedNST cohort, as tumour types (primary, relapsed, etc) were not known.

### Transcriptome data processing

RNA-seq reads from TCGA and TARGET were aligned to human genome 38 (hg38) as described on the GDC website (https://docs.gdc.cancer.gov/Data/Bioinformatics_Pipelines/Expression_mRNA_Pipeline). We used RSEM output for the TCGA and TARGET from Toil (75). The CBTN raw RNA-seq reads were aligned to hg38 using the STAR v2.5.2b (76) and quantified using RSEM v1.2.28 (77) (detailed workflow can be accessed under https://github.com/kids-first/kf-RNA-seq-workflow). The ICGC data were aligned and quantified using STAR v2.7.6a and RSEM v1.2.28, respectively. With the exception of the ICGC, all data processing was performed on the CAVATICA data analysis platform (https://cavatica.sbgenomics.com/). For immune inference, we used transcripts per million (TPM), as per instructions (78). For gene set enrichment scores and consensus clustering, we used log_2_ transformed TPM values corrected for batch effects (data source) using ComBat function from the sva R package. To adjust for cancer type differences, we normalized log_2_ transformed batch-corrected TPM values using median expression in each cancer type.

### Variant and fusion calling

For the TCGA and TARGET, variant calls were obtained from whole exome sequencing (WES) while the ICGC and CBTN variant calls were from whole genome sequencing (WGS). For the TCGA and TARGET, we restricted our analyses to those with MuTect2 calls available on the GDC (details available at https://docs.gdc.cancer.gov/Data/Bioinformatics_Pipelines/DNA_Seq_Variant_Calling_Pipeline). For the ICGC, we used SNV and Indels calls as previously described (23). CBTN WGS data were processed using MuTect2 and Strelka2 (detailed workflows at https://github.com/kids-first/kf-somatic-workflow) and only overlapping calls were used for downstream analysis. For all datasets, we excluded somatic variants with less than 3% variant allele frequency. The Arriba workflow was used for gene fusion calling on CBTN and ICGC datasets (https://github.com/suhrig/arriba). Detailed workflows are available at https://github.com/DKFZ-ODCF/RNAseqWorkflow and https://github.com/kids-first/kf-rnaseq-workflow.

### TMB and oncogenic pathways

To ensure all variants in our study were covered across WES and WGS datasets, we generated a common region list by intersecting bedfiles used in exome experiments with 50 base pairs padding. The exome kits included: Agilent Custom V2 Exome Bait (TCGA and TARGET), Agilent SureSelect All Exon 38 Mb V2 (TCGA), Agilent SureSelect All Exon 50 Mb (TCGA), SeqCap EZ Exome Probes v3.0 (TCGA), SeqCap EZ Human Exome Library v2.0 (TARGET), SeqCap EZ HGSC VCRome 2.1 (TCGA). The final common bedfile consisted of 30,028,393 base pairs. We then used this region list to subset the WGS variant calls. We included non-synonymous coding SNV and Indels in the TMB calculations.

For molecular pathways and their associations with immune clusters, we identified samples with at least one alteration in genes involved in ten TCGA oncogenic pathways (63). The Receptor Tyrosine Kinase (RTK) pathway contained 71 affected genes with a total of 419 alterations in 274 samples that encompassed 71% of pedLGG. Twenty six genes in the PI3-kinase pathway were altered in 74 pedNST samples, 32% of which were pedHGG. The Wnt pathway was disrupted in 70 samples, of which 36% and 20% were MB and CPH. We found 155 alterations in 51 genes involved in the Notch signaling pathway across 66 samples, primarily MB (45%) and pedHGG (23%). Cancer entities with alterations in 32 genes involved in the HIPPO pathway included NBL (25%), pedHGG (22%) and MB (20%). Five core genes involved in the TP53 pathway accounted for 56 mutations across 51 pedNST samples. At least one of 10 cell cycle genes were mutated in 16 pedNST samples, primarily in pedHGG (56%). Acknowledging that *MYCN* amplifications were not included in this analysis, we found

26 alterations in 10 genes related to MYC signalling in 15 samples. Nine samples showed alterations in the TGF-β pathway that consisted of 7 altered genes. NRF2 pathway contained three genes (*KEAP1*, *CUL3* and *NFE2L2*) with 7 mutations across 6 samples. We studied the effects of somatic alterations in the Mismatch Repair (MMR) pathway on immune microenvironment by comparing samples with at least one somatic alteration in *MLH1*, *MSH2*, *MSH6*, *PMS2*, *POLE* or *POLD1* to those without any alterations.

### Immune infiltration and *in silico* simulations

We used ESTIMATE with default parameters to measure overall immune infiltration (https://bioinformatics.mdanderson.org/public-software/estimate/) (26). We performed *in silico* simulations using bulk RNA-seq data from PDX models derived from MB and RNA-seq data of four immune cell-types (CD8+ T-, CD4+ T-, B- cell and NK cells) downloaded from ENCODE portal (https://www.encodeproject.org, experiment IDs: ENCSR861QKF (CD8+ T cells), ENCSR463JBR (CD4+ T cells), ENCSR449GLL (B cells), ENCSR357XTU (NK cells)). For each simulation, an equal percentage of reads from four immune cell-types was randomly sampled using SAMtools v1.9, converted to fastq files with BEDTools v2.27.1 and then concatenated together with randomly sampled reads from PDX RNA-seq data to a total of 100 million reads. The generated pseudo-samples were then processed with Kallisto and Sleuth (79, 80) (108, 109) and TPM values were used as input for the ESTIMATE.

### Immune microenvironment analysis

We used TIMER2 (78) web interface (http://timer.cistrome.org/) for comprehensive analysis of immune-cell composition using six computational tools, CIBERSORT (81), EPIC (82), QUANTISEQ (83), MCPCOUNTER (84), TIMER (85), XCELL (86). We used the CRI-iAtlas Shiny app (https://isb-cgc.shinyapps.io/iatlas/) to cluster the pedNST samples with the immune subtype classifier previously published for adult cancers (14). To estimate immune cell composition using ICGC methylation array data, we used EPIDISH R package (45) with default parameters using reference methylation signatures as previously reported (87).

### Development of immune cell-type specific gene sets and consensus clustering

To identify genes specific to the immune system, we leveraged four data sources: 1) We obtained median gene expression for 18 purified cell-types, as previously reported (13). We found 9,897 genes were expressed > 75^th^ percentile of normalized expression (median centered and Median Absolute Deviation (MAD) scaled) in at least one cell-type within this dataset. We selected 1,958 genes with ≥ 2 MAD difference between immune and non-immune cell populations. 2) We downloaded the human protein atlas data v20.1 (https://www.proteinatlas.org/about/download). The Human Protein Atlas consisted of single-cell data from 51 cell populations (40). 15,302 genes were expressed > 75^th^ percentile in at least one cell population. 3) In the Human Protein Atlas and across 37 tissues, 11,069 genes were annotated as specific to at least one tissue, determined as normalized expression ≥ 1 (median normalized expression 33.9). We found 7,206 genes specific to only one tissue, 1,257 of which were specific to blood, lymphoid tissue or bone marrow. 4) We included the ESTIMATE immune signature consisting of 141 genes (26). In total, we found 3,041 unique genes across four data sources with evidence of specificity to immune cell populations.

Next, we sought to identify genes that may be expressed in pediatric cancer and other non-immune cells. We compiled a pediatric cancer geneset using three data sources. 1) We used protein-coding gene expression data from 79 PDX models. We found 13,788 genes expressed > 75^th^ percentile in at least one PDX. 2) We used data from 22 cell lines collected by CBTN. These consisted of one EPN, 18 pedHGG and 3 MB. Across these cell lines, 11,195 protein-coding genes were expressed > 75^th^ percentile in at least one cell line. 3) We leveraged published single-cell datasets from pediatric cancers, NBL (16) and EPN (18, 19) and used original cell annotations as reported by authors. As single-cell datasets are generally sparse, we focused our analysis to genes with maximum expression > 75^th^ percentile in at least one cell population in each dataset. We then determined non-immune genes as those with ≥ 2 MAD difference between non-immune and immune cell populations in each dataset. This analysis resulted in 3,036, 12,109 and 5,040 non-immune genes from Jansky, Gojo and Gillen datasets, respectively. Intersecting the immune and non-immune genes resulted in 791 immune-specific genes.

To accurately assign the identified genes to immune cell-types, we compiled genesets from seven sources and conducted a consensus approach to assign genes to specific immune cell-types. For this analysis, we aggregated immune subtypes (e.g. Tregs) into the following major immune cell-types: T, B, NK cells, dendritic cells, monocytes, macrophages, granulocytes and myeloid cells. These sources included gene annotations from five immune deconvolution tools, MCPCOUNTER, QUANTISEQ, CIBERSORT, EPIC and TIMER. We used all 264 genesets that were used to develop XCELL. We used two additional sources: 1) scaled average counts of 7,172 genes specific to 28 purified immune cell types derived from 416 healthy donors and patients with immune diseases (Immunexut) (41). We chose 3,896 genes with scaled count > 75^th^ percentile in at least one cell-type, of which 2,555 genes had ≥ 2 MAD difference between one cell-type compared to all others. 2) The human protein atlas annotated 5,934 genes as blood cell specific (42), determined as genes with normalized expression ≥ 1 in at least one cell-type. We chose 4,902 genes that were specific to one immune cell-type. Our consensus analysis revealed 216 genes that were assigned to one specific immune cell-type in at least two independent sources. We excluded myeloid-cell geneset due to the low number of genes (n = 6) that may lead to inaccurate enrichment scores. To calculate enrichment scores, we applied single-sample geneset enrichment analysis (ssGSEA) from GSVA R package (88) to batch-corrected log_2_-transformed TPM values across pedNST. We then applied consensus clustering to the scaled (median centered and MAD scaled) immune cell-type enrichment scores using the ConsensusClusterPlus R package (89). We performed k-means clustering based on Euclidean distance with 200 subsamples using 80% of samples and 100% of features for 2 to 8 clusters. This analysis revealed four major immune clusters across pedNST.

For further delineate types of infiltrating T cells, we derived enrichment scores of 40 T-cell subtypes using top 50 signature genes derived from recent pan-cancer analysis of tumour-infiltrating lymphocytes (90). We focused this analysis on Pediatric Inflamed to avoid overestimation as a result of low overall immune infiltration in other clusters. We scaled enrichment scores within each T-cell subtype across Pediatric Inflamed and performed consensus clustering with parameters as previously described. For myeloid-cell subtypes, we used 13 myeloid signatures derived from single-cell analysis of adult cancer patients (91). We chose 9 gene signatures consisting of genes with ≥ 1.5 fold change in expression between any given myeloid cell-type and other clusters. We scaled enrichment scores from myeloid gene signatures within each myeloid cell subtype across Myeloid Predominant and performed consensus clustering with parameters as previously described. For microglia, we used signatures from two previous reports and derived enrichment scores in the pedCNS subset of Myeloid Predominant using ssGSEA (52, 92).

### Immunohistochemistry

Formalin-fixed, paraffin-embedded TMA sections were analyzed for CD4, CD8 and CD19 expression. The IHC staining was performed using the Ventana Discovery platform. IHC was optimized and performed with CD4 (Abcam Ab183685), CD8 (Leica NCL-L-CD8-4B11) and CD19 (e-Bioscience 14-0194) with dilutions of 1:500, 1:100 and 1:500 respectively. In brief, tissue sections were incubated in Tris EDTA buffer (cell conditioning 1; CC1 standard) at 95C for 1 hour to retrieve antigenicity, followed by incubation with the respective primary antibody for 1 hour. Bound primary antibodies were incubated with the respective secondary antibodies (Jackson Laboratories) with 1:500 dilution, followed by Ultramap HRP and Chromomap DAB detection. For staining optimization and to control for staining specificity, normal tonsil was used as control. Intensity scoring was done on a common four-point scale. Descriptively, 0 represents no staining, 1 represents low but detectable degree of staining, 2 represents clearly positive staining, and 3 represents strong expression. Expression was quantified as H-score, the product of staining intensity and percentage of stained cells. The TMAs used in the study were obtained from the Children’s Oncology Group (COG) and contains 9 ATRT, 13 pedHGG, 20 EPN, 64 MB and 33 NBL cases, all of which are represented in duplicate cores.

To study protein levels of TIM3 and LAG3 in NBL, we purchased two serial TMA slides consisting of 26 NBL cases all in duplicate cores with 10 cores of normal peripheral nerve tissue (Biomax, NB642a). Briefly, TMA sections were dewaxed, rehydrated through an ethanol series to water and endogenous peroxidases were blocked using 3% H_2_O_2_ in PBS for 15 min at room temperature. Antigen retrieval was performed using 10 mM sodium citrate pH 6.0. Primary antibodies were incubated for 45 min at room temperature (TIM3 CST45208 1:150, LAG3 ab40466 1:200) followed by several washes with PBS. Secondary antibodies were incubated for 30 min at room temperature (BA-1000 and BA-9200 at 1:500 dilutions Vectors Labs). ABC kit (PK-6100, Vectors Labs) was applied for 25 min with DAB for 4 min (SK-4100) followed by PBS washes. Stained slides were scanned using a Nanozoomer 2.0HT (Hamamatsu Photonics) at 20x or 40x.

### TIL Analysis on H&E Whole Slide Images

Computational tumour-infiltrating lymphocyte (TIL) analysis was performed on digitally scanned hematoxylin and eosin (H&E)-stained whole-slide images (WSI) from the Children’s Brain Tumour Network (CBTN) as previously described (44). Briefly, a pre-trained Inception-V4 deep learning model was applied to each WSI, returning a model probability of TIL presence within 50×50 micron patches tiling across the entire WSI. Image patches that did not contain tissue were filtered out using a color standard deviation threshold of 18 (computed on an 8-bit [0,255] color scale) and a mean TIL score was computed as the average TIL probability on the remaining patches for each WSI. Upon manual review, WSI with poor scanning quality and/or with a high false positive rate of TIL detection were removed from the analysis, including medulloblastoma and other tumours arising in the cerebellum/posterior fossa. One outlier sample was removed (Grubbs test).

### HLA-typing and prediction of neoantigens

We used the Optitype tool available on the CAVATICA platform to determine HLA-A, B and C types from the TARGET and CBTN RNA-seq datasets (60). To predict neoantigens from gene mutations, we used Mutant Peptide eXtractor and Informer (MuPeXI) tool using MuTect2 variant caller as input (61). We predicted 8-11mer peptides for all HLA-types. We determined strong or weak binding peptide-MHC complexes as previously described (percentile rank ≤ 0.5 for strong binding, > 0.5 and ≤ 2 for weak binding peptides) (62). We included peptides predicted from mutations that overlapped between MuTect2 and Strelka2.

### T- and B- cell repertoire analysis

To recover T- and B- cell clonotypes, we used MiXCR v2.1.12 (65) with default parameters for RNA-seq data processing. We applied the framework iNterpolation/EXTrapolation (iNEXT) to study immune repertoire diversity (93). This method was primarily developed for diversity estimates in ecology and aimed to bring together asymptotic estimates and rarefaction/extrapolation methods of estimation for samples with different sizes. Three most popular diversity indices have already been translated to immune repertoire studies; richness, the total number of species, Shannon index (entropy), which puts moderate weight on abundance and Simpson index, which measures diversity of abundant species. The robustness of these indices is dependent on sample size and could be biased in under-sampled experiments, which is a known problem in biodiversity studies and attempts have been reported to reduce such empirical biases (94). Chao et al. provided a more accurate estimation of diversity by integrating slopes obtained from accumulation curves into entropy formula followed by bootstrapping method for variance (94). We applied similar principles on RNA-seq datasets to infer immune diversity. We found that the estimates were severely affected in samples with less than three clonotypes recovered from bulk RNA-seq or in samples with multiple clonotypes of the same clonal fractions and excluded them from our analysis. Note that these samples may be A) samples with low number of immune cells and therefore low number of clonotypes that could not be reflected on shallow RNA-seq data, or B) samples with sufficient immune infiltration, but highly uneven clonal distribution, therefore only highly abundant clonotypes were seen at shallow coverage, yet the true diversity could not be robustly estimated.

For B-cell repertoire, we used constant regions of immunoglobulin heavy chain (IGH C segments) to study the distribution of immunoglobulin isotypes. Due to somatic hypermutation in the B-cell repertoire, we removed sequences with ≤ 2 reads. As B-cell clones originating from a naïve B cell may have mismatches in CDR3 sequences via somatic hypermutation events, we clustered IGH CDR3 octamers allowing for one mismatch, as described previously (67). We considered sequences shorter than 8 amino acids as individual CDR3 sequences. Only samples with > 3 CDR3 sequences were included from this analysis and we used the gini index of inequality as a measure for uneven distribution of B-cell clusters.

### TCR hybrid capture sequencing

To validation the T-cell diversity estimates from RNA-seq, we applied CapTCR-seq (66) hybrid capture protocol to RNA-seq libraries of adult nasopharyngeal carcinoma (n = 33), pediatric samples from the PRecision Oncology For Young peopLE (PROFYLE) program (www.profyle.ca) (n = 10) and ten pediatric GBM samples (ICGC). We compared estimated values of TCRβ richness, Shannon and Simpson diversities from bulk RNA-seq data with observed richness, Shannon and Simpson diversities from CapTCR-seq experiment.

### Statistics and visualization

Statistical tests, p-values and other details are noted in text and figure legends. For tumour subtypes, we used one-sided Fisher’s exact test (alternative = “greater” in R) comparing each subtype to all others within each immune cluster. We used analysis of covariance (ANCOVA) to compare continuous variables such as TMB or age at diagnosis across immune clusters adjusting for cancer entities. We used log-rank tests for Kaplan-Meier analyses. We used Cox proportional hazards models to adjust for cancer entities and gender in multivariable analyses. We used Student’s t-test to compare scaled values across groups and rank sum test to compare values with non-normal distribution. We applied the Cochran-Mantel-Haenszel (CMH) test to compare samples with mutations in oncogenic pathways across immune clusters controlling for the effects of cancer entities. In all boxplots, boxes show median and IQR and whiskers represent 1.5 times IQR. Boxplots are shown for groups with more than three datapoints. We annotated statistical significance levels as follows: *p < 0.05, **p < 0.01 and ***p < 0.001. We performed all analyses and visualizations in R v4.0. We used Adobe Illustrator v24.0.1 for aesthetic edits and figure alignments.

### Source code and data availability

CBTN raw RNA and WGS data, variant and fusion calls are available upon request at kidsfirstdrc.org. TCGA and TARGET data can be accessed upon request through GDC (https://gdc.cancer.gov/). Gene expression matrices, clinical metadata, T- and B- clonotype files, HLA types, predicted neopeptides, variant and fusion datasets and deconvolution outputs are available upon free registration at https://cavatica.sbgenomics.com/u/pughlab/immpedcan. Custom codes and input data are available at https://github.com/pughlab/immunogenomics_pedNST. Main figures can be reproduced via CodeOcean capsule: https://doi.org/10.24433/CO.0339991.v1

## Declaration of Interests

J.N.P is an employee of Genentech, Inc. (Stock and Other Ownership Interests: F. Hoffmann-La Roche AG).

## Acknowledgements

This research was supported by a Terry Fox Research Institute New Investigator Award (TJP), the Gabriella Miller’s Kids First Data Resource Center (NIH common funds), the Dragon Masters Foundation and the ITCC-P4 consortium (Innovative Medicines Initiative2 Joint Undertaking, under grant agreement No. 116064). Additional funding support came from the PedBrain Tumour Project contributing to the International Cancer Genome Consortium (ICGC) funded by German Cancer Aid (109252) and the German Federal Ministry of Education and Research (BMBF, NGFN^plus^ #01GS0883). AN was supported by a Princess Margaret Post-doctoral Fellowship. PB is funded by the DKFZ International PhD Program (Annemarie Poustka fellowship). TJP is supported by the Canada Research Chair in Translational Genomics, a Senior Investigator Award from the Ontario Institute for Cancer Research, and the Princess Margaret Cancer Foundation Gattuso-Slaight Personalized Cancer Medicine Fund. DAO is supported by NHLBI R38 HL143613, NCI T32 CA009140, and the Parker Institute for Cancer Immunotherapy. We thank Ben Wang and Andrew Elia of Tumour Immunotherapy Program at Princess Margaret Cancer Centre for performing immunohistochemistry experiments (pm-tumourimmunotherapy.ca). We thank Wei Xu, Jingyue Huang and Osvaldo Espin-Garcia (Dana Lana School of Public Health, University of Toronto) for statistical assistance. The data and materials used for the immune repertoire analysis were made available by Nada Jabado, and the PRecision Oncology For Young peopLE (PROFYLE) program and through funds from the Terry Fox Research Institute and all other funders supporting PROFYLE. We thank Pierre Antoine (Princess Margaret Cancer Centre), Angela Bik-Yu Hui (Stanford Cancer Institute) and Fei-Fei Liu (Princess Margaret Cancer Centre) for providing nasopharyngeal carcinoma samples for immune diversity measurements. We also thank the staff of the Princess Margaret Genomics Centre (www.pmgenomics.ca) and Bioinformatics and High-Performance Computing Core for their expertise in generating the immune repertoire sequencing data used in this study.

## Supplemental figures

**Figure S1. Characterization of the CBTN dataset. A-B)** Heatmaps showing hierarchical clustering of CBTN samples using top 1000 variable genes before (A) and after (B) removing samples that pathological annotations did not match their transcriptional clusters. **C)** Sankey plot showing the CBTN samples and their annotations included in this study. **D-E)** Heatmaps showing hierarchical clustering using top 1000 variable genes for the CBTN MB (D) and ATRT (E) samples to identify tumour subgroups.

**Figure S2. Assessment of ESTIMATE performance using *in silico* simulations.** Scatter plot and non-linear regression line showing correlation between the ESTIMATE Immune score and immune read percentage in simulated PDX samples with increasing percentage of immune reads.

**Figure S3. Immunohistochemistry confirms variable immune infiltration in pedNST. A)** Boxplots showing H-scores for anti-CD8, CD4 and CD19 staining across an independent cohort of pedNST. **B)** Representative images for immunohistochemistry. In all boxplots, boxes show median and IQR and whiskers represent 1.5 times IQR.

**Figure S4. Analysis of immune deconvolution tools reveals discordance in immune inference. A-F)** Heatmaps representing immune cell-type estimates derived from CIBERSORT (A), EPIC (B), TIMER (C), MCPCOUNTER (D), XCELL (E) and QUANTISEQ (F) across PDX and pedNST. Dendrograms show hierarchical clustering between cell-types (rows) and samples (columns). **G)** Barplots obtained from TIMER2 comparing estimates of 8 immune cell-types (columns) derived from six immune deconvolution tools (rows) in 10 PDX MB samples (bars).

**Figure S5. Adult tumour microenvironment clusters and tumour purity analysis confirm characteristics of immune clusters. A)** Heatmap illustrating fraction of samples in each CRI-iAtlas cluster across immune clusters in pedNST. Barplot shows total number of samples in each CRI-iAtlas cluster. **B)** Boxplot depicting tumour purity scores for 156 ICGC samples inferred from copy number estimates (ACEseq). Boxes show median and IQR and whiskers represent 1.5 times IQR. **C)** Representative plots showing B allele frequencies (BAF) and copy numbers (TCN) in each immune cluster.

**Figure S6. Segmentation analysis of H&E images and methylation-based deconvolution analysis validate transcriptional immune clusters. A-B)** Boxplots showing average TIL score based on segmentation analysis of pathological images across cancer entities (A) and tumour sites (B) using H&E images from CBTN. **C)** Boxplots showing average TIL score based on segmentation analysis of pathological images in samples with > 5% immune read percentage (left panel), in samples collected from cerebral hemisphere (right panel). **D)** Ridge plots illustrating z-score distributions for immune cell-types as estimated by methylation-based deconvolution analysis across immune clusters. Pairwise two-sided Student’s t-test with Bonferroni correction, **p < 0.01, ***p < 0.001. Significance levels for T cells are shown compared to Pediatric Inflamed. Significance levels for all other cell-types are shown compared to Myeloid Predominant. In all boxplots, boxes show median and IQR and whiskers represent 1.5 times IQR.

**Figure S7. Distribution of clinical parameters across cancer entities. A-B)** Barplots showing fraction of gender (A) and ethnicity (B) across cancer entities in pedNST. **C)** Boxplot depicting age at diagnosis (in years) across cancer entities in pedNST. Boxes show median and IQR and whiskers represent 1.5 times IQR.

**Figure S8.** Profiling immune checkpoint genes reveals a subgroup of neuroblastoma with elevated *HAVCR2 and* high *LAG3* expression in Immune Excluded. A) Boxplot showing expression of five targetable immune checkpoint genes in 79 PDX models derived from ATRT, EPN, MB, NBL and pedHGG. **B)** Boxplots showing *LAG3* gene expression in the Immune Excluded cluster across all cancer types. **C)** Heatmap showing expression of five immune checkpoint genes in two independent datasets, TARGET (top) and Kids First NBL (bottom). **D)** Protein levels of TIM3 and LAG3 in an independent NBL tissue microarray. Left panel shows staining H-scores obtained from a pathologist’s review. Representative images of high TIM3/low LAG3 and low TIM3/low LAG3 are shown on the right. In all boxplots, boxes show median and IQR and whiskers represent 1.5 times IQR.

**Figure S9. HLA class I frequencies in pedNST.** Barplots showing frequency of HLA-A, B and C across pedNST.

**Figure S10. Distribution of predicted peptides across cancer entities and immune clusters. A-B)** Boxplots showing predicted strong (A) and weak (B) binding peptides across cancer entities in pedNST. **C-D)** Boxplots depicting predicted weak binding peptides in the pedNST (C) and pedHGG (D) samples across immune clusters. Two-sided rank sum test, *p < 0.05. In all boxplots, boxes show median and IQR and whiskers represent 1.5 times IQR.

**Figure S11. Validation of T- and B- cell repertoire analysis and distributions across cancer entities. A-B)** Scatterplots depicting linear correlation between TCRβ Simpson diversity (A) or richness (B) estimated from RNA-seq and measured by capturing TCR sequences from the same RNA-seq libraries in adult and pediatric cancer samples**. C)** TCRβ estimated Shannon diversity across cancer entities in pedNST ordered by median. **D)** Barplots showing total number of immunoglobulin isotypes normalized by number of samples in each cancer type (top barplot) and isotype fractions (stacked barplot) in each cancer entity. **E)** The gini index for B-cell repertoire across cancer entities in pedNST ordered by median. Circle plots show two CPH samples with gini indices of 0.74 (top) and 0.22 (bottom). Each circle is one B-cell clone and circle diameters are proportional to immunoglobulin reads. Blue circles denoted clusters of highly similar immunoglobulin sequences.

**Figure S12. Analysis of immune-cell subtypes in Pediatric Inflamed and Myeloid Predominant provides insights into mechanisms of immunosuppression. A)** Heatmap showing scaled gene set enrichment scores for 40 T-cell subtypes as annotated in (90) in the Pediatric Inflamed (Tn: Naïve T cells, Th: T helper, ISG: Interferon-stimulated genes, Tm: memory T cells, Trm: Tissue-resident memory T cells, Tem: effector memory T cells, Temra: effector memory re-expressing CD45RA T cells, Treg: regulatory T cells, Tfh: follicular helper T cells, Th17: T helper 17, Tex: exhausted T cells, KIR: Killer immunoglobulin-like receptor). **B)** Boxplots showing log-transformed TCRβ estimated Shannon diversity (left) and TMB (right) across T-cell groups (TG). Pairwise two-sided Student’s t-test, *p < 0.05, **p < 0.01 and ***p < 0.001 (left). Two-sided rank sum test, *p < 0.05 (right). **C)** Boxplots showing differences in immune-cell infiltration, as determined in Fig1C, across T-cell groups (TG). Pairwise two-sided Student’s t-test, *p < 0.05, **p < 0.01 and ***p < 0.001. **D)** Heatmap illustrating median tumour-type normalized expression of selected genes in TG2 and TG5. **E)** Heatmap depicting scaled gene set enrichment scores for 9 myeloid-cell subtypes as annotated in (91, 95) (DC: Dendritic cell, Mono: Monocyte, Mac/Macro: Macrophage, TAM: Tumour-associated macrophage). **F)** Heatmap showing scaled gene set enrichment scores for microglia (52, 92) in the pedCNS subset of Myeloid Predominant. **G)** Heatmap illustrating median tumour-type normalized expression of selected genes in MG1, MG3 and MG5. In all boxplots, boxes show median and IQR and whiskers represent 1.5 times IQR.

## Supplemental Note. Related to FigS12

### Lymphoid and Myeloid subtypes provide insights into mechanisms of immune evasion

To further characterize T-cell subtypes in the Pediatric Inflamed cluster with high levels of lymphocyte infiltration, we obtained 40 gene signatures of CD4+ and CD8+ T-cell subpopulations from the single-cell profiling of tumour-infiltrating lymphocytes across 21 adult cancers (90). Consensus clustering revealed five distinct T-cell groups (TGs) within Pediatric Inflamed with notable patterns related broadly to naïve, exhausted and memory T-cell populations (FigS12A). Nine NBL samples were characterized by elevated enrichment of naïve T cells (TG2). TG3 consisted of 6 samples with enrichment of Naive and NME1+ T cell signatures while depletion of other T cell subtypes. TG4 (n = 30) showed intermediate enrichment for T-cell signatures and included CPH, NFB, MNG and MYC-like ATRT. TG5 was a group of 10 samples with highest enrichment of T-cell signatures related to memory (Tm), effector memory (Tem), T follicular helper/helper (Tfh/h), Tregs and exhausted (Tex) T cells. These data identify subset of samples within Pediatric Inflamed that are infiltrated with naive (TG2/TG3) or differentiated (TG4/TG5) T cells.

In Pediatric Inflamed, Cox proportional hazards model adjusting for gender and cancer type revealed that TG5, highly infiltrated with differentiated T cells, had worse PFS compared to TG1 (HR = 9, 95% CI: 1.5-54, p = 0.02). TG5 also showed lower T-cell diversity compared to TG2 and TG3 that harbored high enrichment of T cell naïve and proliferating NME1+ T-cell signatures (FigS12A) suggesting T-cell clonal expansion in TG5 (p = 0.0002 and 0.04, FigS12B, left). While TG2 harbored significantly higher TMB compared to TG1 (p = 0.02), TG5 did not show any significant difference in TMB compared to other groups (FigS12B, right). Compared to TG2, TG5 harbored higher infiltration of T, NK and myeloid cell infiltration suggesting antigen recognition by myeloid cells followed by clonal expansion and T-cell differentiation (p < 0.008, FigS12C). TG5 showed higher expression levels of HLA class I genes (*HLA-A*, *HLA-B*, *HLA-C*), inhibitory cytokines and chemokines (*CXCL8, CXCL1, IL6*), genes encoding transcription factor AP-1 subunits involved in IFN-γ pathway (*FOS, FOSB, JUN, JUNB, ATF3*), markers of T-cell activation (*CD86*, *CD69*) and Toll-like receptors (*TLR2*, *TLR4*) (FigS12D). Conversely, TG2 showed higher levels of B-cell marker genes (*CD19*, *CD22*), immune checkpoint gene (*BTLA*) and Fc receptors (*FCER2*, *FCRL2*) (FigS12D) suggesting a B-cell mediated mechanism of T-cell suppression. These results suggest two distinct mechanisms of immunosuppression in Pediatric Inflamed, characterized by suppression of T-cell effector functions (TG5) and inhibition of T-cell activation and differentiation (TG2).

To delineate myeloid-cell subtypes in Myeloid Predominant, we performed consensus clustering using recently described signatures for tumour-infiltrating myeloid cells (91, 95) (methods). MG1 consisted of 22 samples with high enrichment of mast cells, *CD1C*-expressing conventional dendritic cells type 2 (cDC2), monocytes and macrophages (FigS12E). MG2 (n = 89) showed low enrichment of dendritic cells while high levels of *PLTP*-expressing macrophages. MG3 (n = 70) was characterized by elevated enrichment of *LILRA4*-expressing plasmacytoid and *BATF3*-expressing conventional dendritic cells (pDC and cDC1) and tumour-associated macrophages (TAM). MG4 (n = 75) showed depletion of mast and macrophage signatures with moderate enrichment of monocytes and cDC1 signatures. MG5 (n = 23) had the lowest enrichment of all myeloid signatures (FigS12E). Scoring microglia-specific gene signatures (52, 92) in CNS tumours showed pedLGG samples had the highest levels of microglia geneset enrichment predominantly in MG3 (FigS12F). These findings reveal two prominent subgroups within Myeloid Predominant, characterized by high levels of Mast cells, macrophages and monocytes (MG1) and high levels of pDC and TAMs (MG3).

Cox proportional hazards model adjusting for gender and cohort revealed that low Mast/Macro groups (MG3, MG4 and MG5) had a trend toward improved PFS compared to MG1 (HR = 0.4, 0.38 and 0.37, 95% CI: 0.14-1.14, 0.14-1.02 and 0.12-1.12, p = 0.09, 0.05 and 0.08, respectively). To identify potential mechanisms underlying MGs, we performed differential gene expression analysis comparing MG1, MG3 and MG5 controlling for cancer types. In MG1, genes associated with cytotoxic T cells (*FGFBP2*) and leukocyte trafficking (*TSPAN8*) (96) were expressed at higher levels compared to MG3 and MG5 (FigS12G). We found high expression of *PENK*, encoding an opioid receptor proenkephalin and a precursor for neuropeptides suggesting crosstalk between immune and neural cells (97) (FigS12G). *ANGPTL7*, encoding angiopoietin-like protein 7, that regulates angiogenesis (98) and *F2RL2,* encoding proteinase-activated receptor 3 (PAR3), that may trigger myeloid infiltration through coagulation initiation (99) were expressed at higher levels in MG1. These were accompanied by high expression of collagens (*COL14A1, COL23A1*) that are known components of the extracellular matrix that impact T-cell infiltration (100, 101). Additionally, we found *IL34*, a brain-specific ligand for CSF1R, was expressed at higher levels in MG1 suggesting a role in monocyte-macrophage differentiation in MG1 (102, 103). In MG3, genes involved in endothelial regulation (*ESM1*, *APLN*, *ALCAM*) were expressed at higher levels compared to MG1 and MG5 (Fig6G). *EDIL3* and *POSTN* encoding ligands for αVβ3 integrin (DEL-1 and periostin) were expressed at higher levels in MG3, the latter of which can recruit tumour-associated macrophages (TAM) in adult glioblastoma (104, 105). Genes encoding the αVβ3 integrin (*ITGAV*, *ITGB3*) were also expressed at higher levels in MG3, suggesting periostin as a regulator of TAM infiltration. Interestingly, Histamine receptor H1 (*HRH1*) and *HNMT*, a gene involved in histamine metabolism were both expressed at higher levels in MG3 (FigS12G) suggesting a mechanism of immune evasion and T-cell dysfunction (106). These findings suggest cellular interactions influencing the myeloid compartment (periostin-integrins, IL34-CSFR1) and immune exclusion and evasion (histamine and collagens) in Myeloid Predominant.

## Supplemental tables

**TableS1. Overview of patient characteristics and analyses.**

**TableS2. Immune cell-type estimates derived from immune deconvolution tools.**

**TableS3. Curated signature genes used to define immune clusters.**

**TableS4. Tumour-infiltrating lymphocyte scores for each sample.**

**TableS5. Cox proportional hazards model for associations of immune clusters with OS.**

**TableS6. Cox proportional hazards model for associations of immune clusters with PFS.**

**TableS7. Results from QuSAGE Pathway analysis across immune clusters.**

**TableS8. Tumour mutation burden values for each sample.**

**TableS9. ANCOVA for total SNV/Mb among immune clusters.**

**TableS10. ANCOVA for total SNV_Indel/Mb among immune clusters.**

**TableS11. ANCOVA for total strong binding neopeptides among immune clusters.**

