## Supplemental Figures for "Transcriptional immunogenomic analysis reveals distinct immunological clusters in pediatric nervous system tumours"

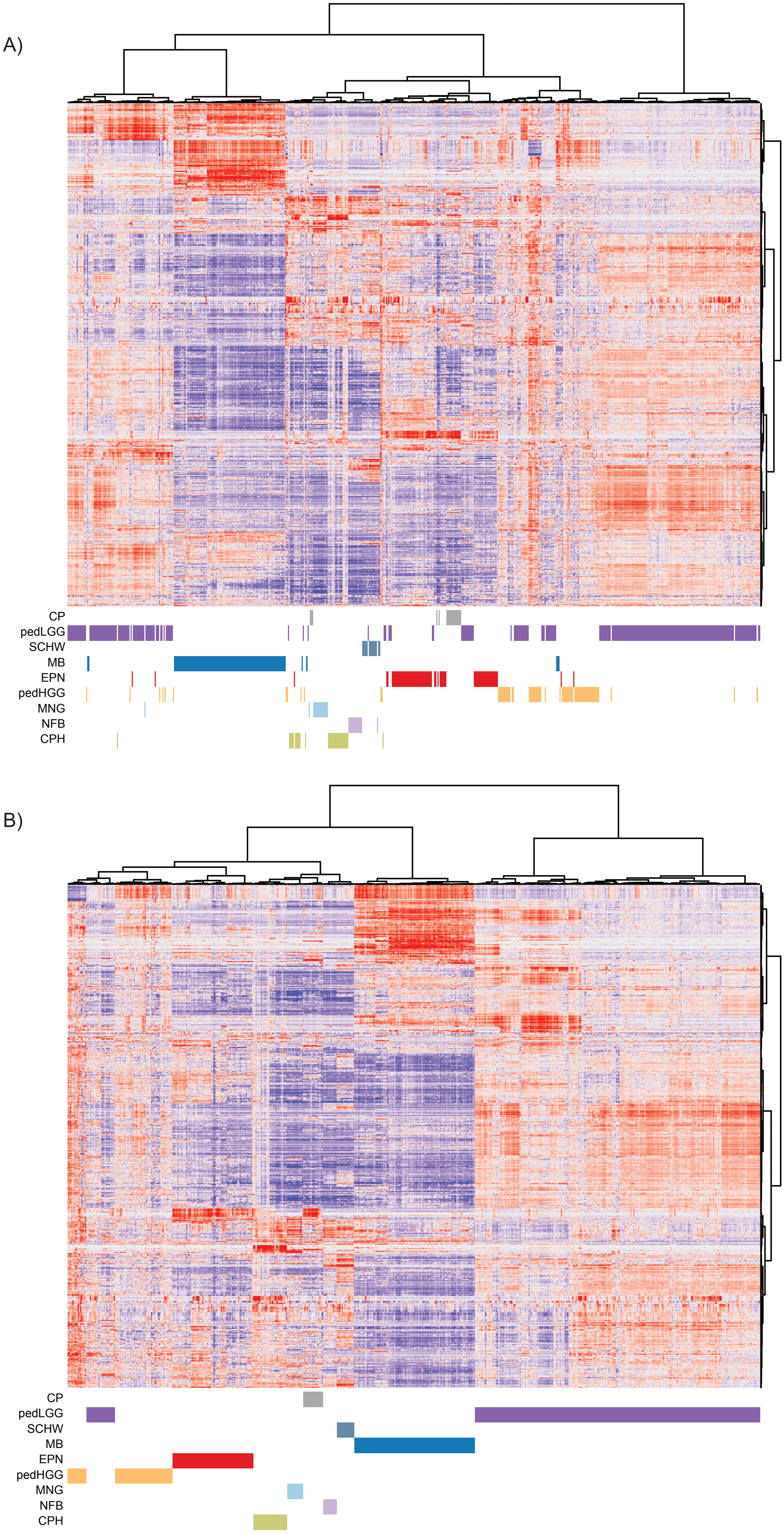
A)
CP
pedLGG
SCHW
MB
EPN
pedHGG
MNG
NFB
CPH
B)
CP
pedLGG
SCHW
MB
EPN
pedHGG
MNG
NFB
CPH

Figure S1

C)

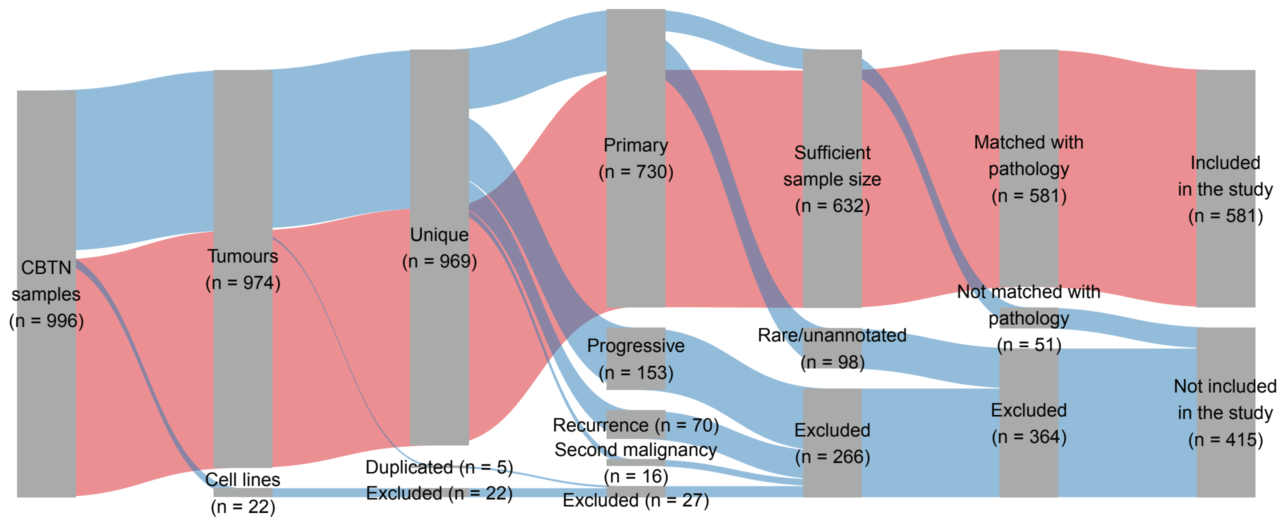

CBTN samples (n = 996)
Tumours (n = 974)
Cell lines (n = 22)
Unique (n = 969)
Duplicated (n = 5)
Excluded (n = 22)
Primary (n = 730)
Progressive (n = 153)
Recurrence (n = 70)
Second malignancy (n = 16)
Excluded (n = 27)
Sufficient sample size (n = 632)
Rare/unannotated (n = 98)
Excluded (n = 266)
Matched with pathology (n = 581)
Not matched with pathology (n = 51)
Excluded (n = 364)
Included in the study (n = 581)
Not included in the study (n = 415)


D)

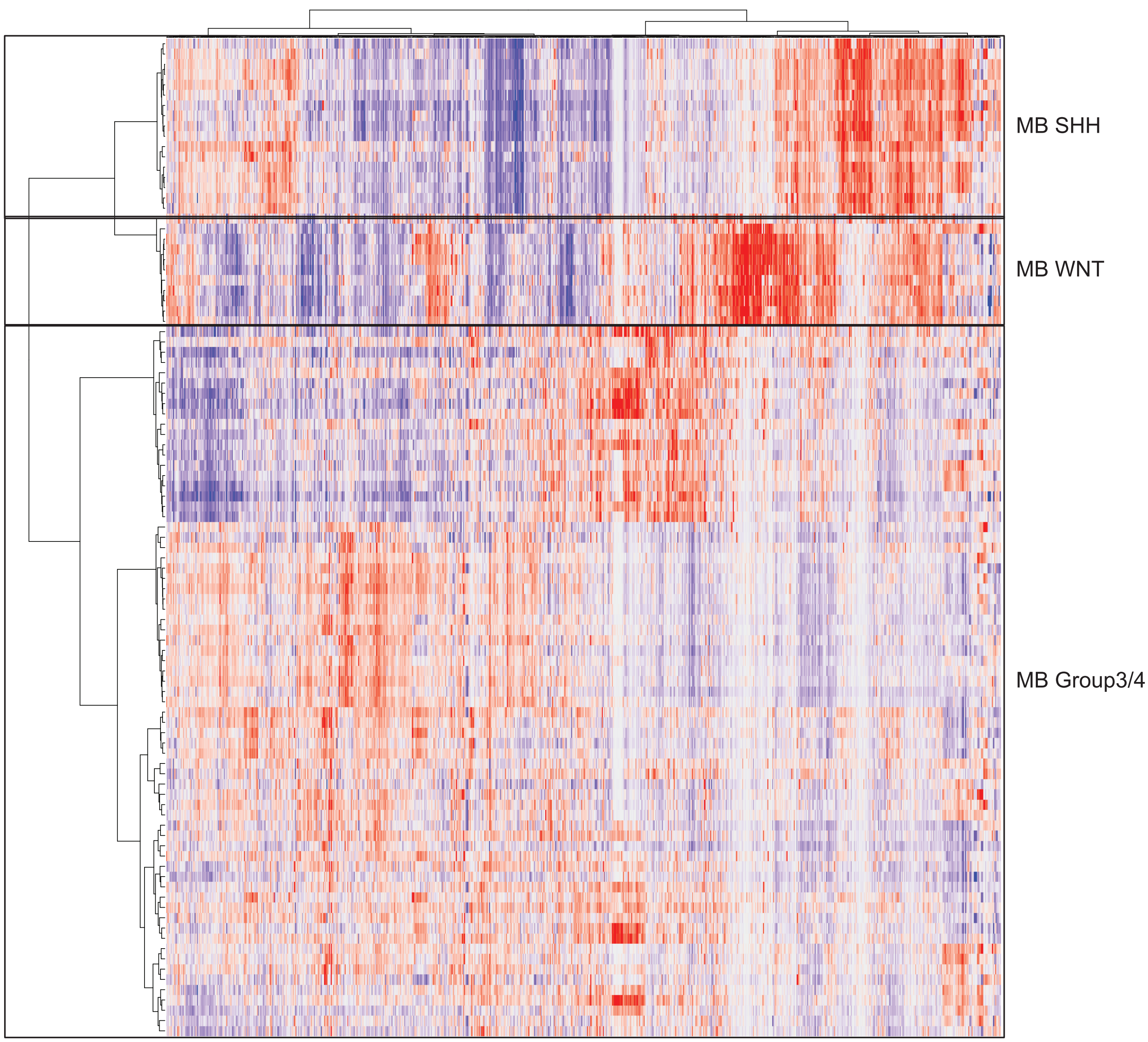

MB SHH
MB WNT
MB Group3/4


Figure S1

E)

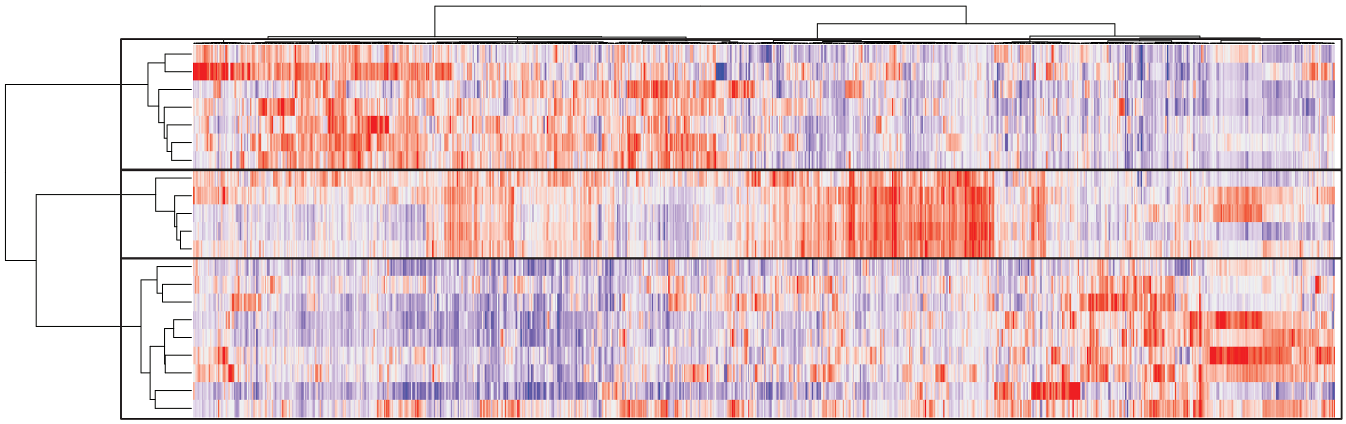

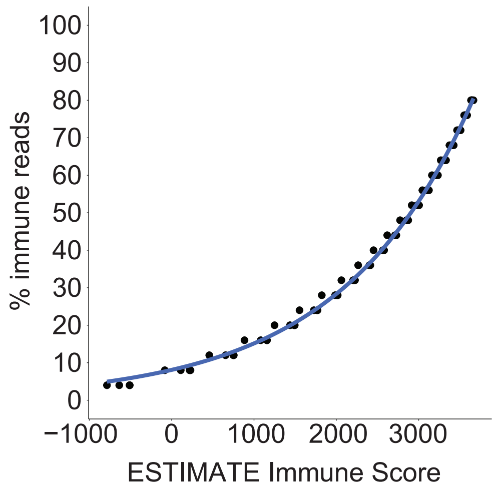

% immune reads
100
90
80
70
60
50
40
30
20
10
0
−1000
0
1000
2000
3000
ESTIMATE Immune Score


Figure S2

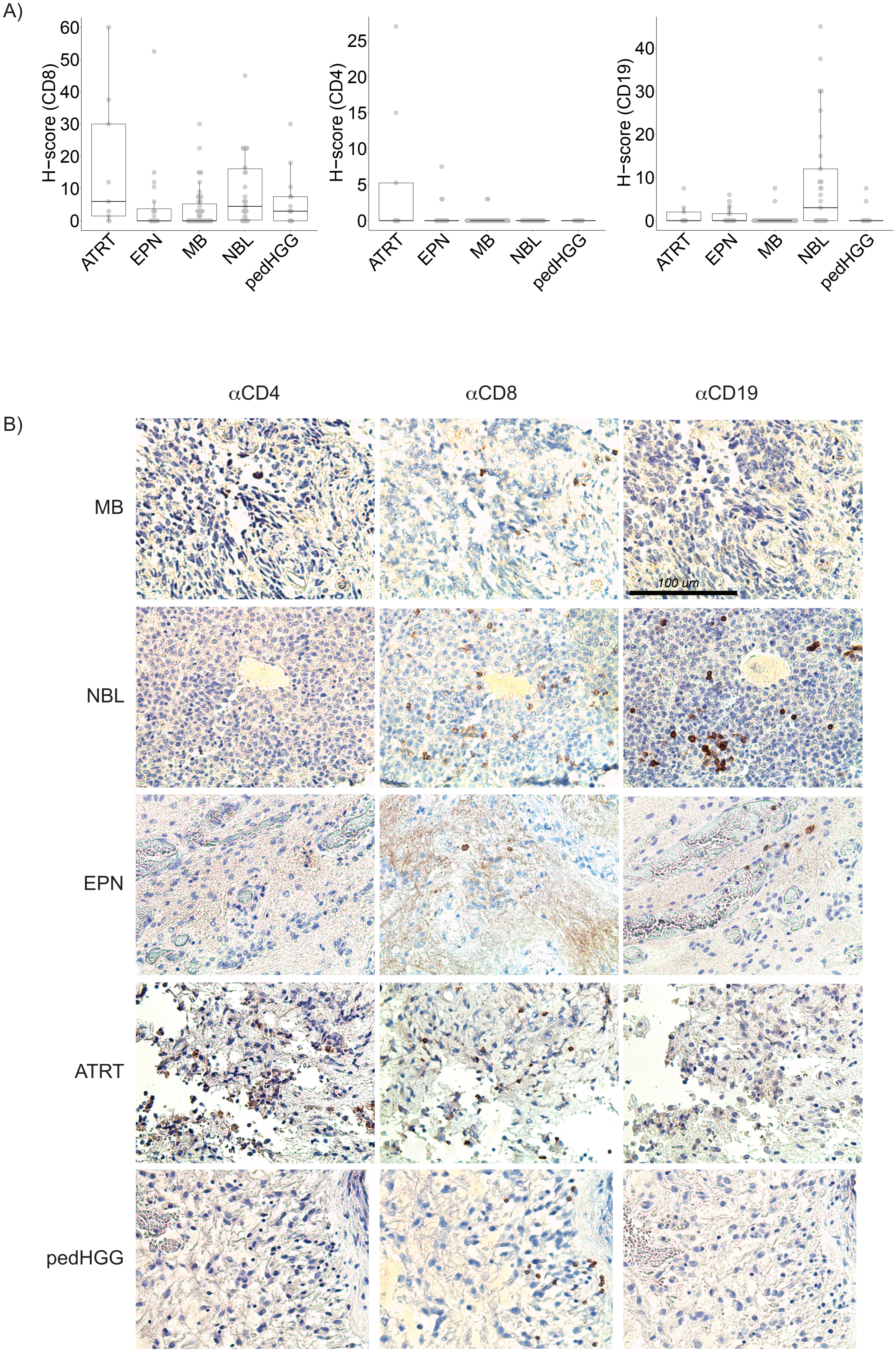
A)
H−score (CD8)
60
50
40
30
20
10
0
ATRT
EPN
MB
NBL
pedHGG
H−score (CD4)
25
20
15
10
5
0
H−score (CD19)
40
30
20
10
B)
αCD4
αCD8
αCD19
MB
NBL
EPN
ATRT
pedHGG
100 um

Figure S3

A)

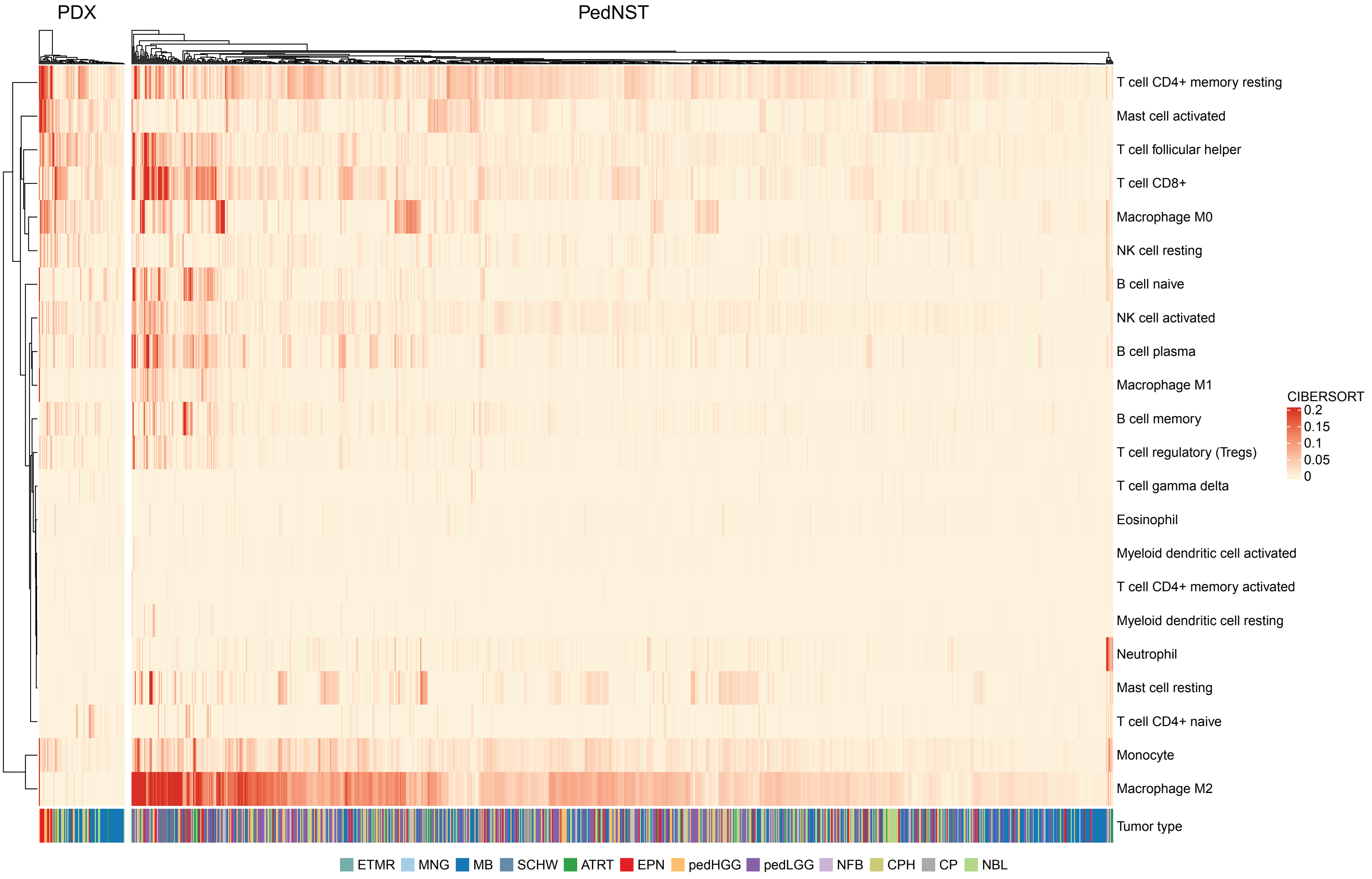
PDX
PedNST
T cell CD4+ memory resting
Mast cell activated
T cell follicular helper
T cell CD8+
Macrophage M0
NK cell resting
B cell naive
NK cell activated
B cell plasma
Macrophage M1
B cell memory
T cell regulatory (Tregs)
T cell gamma delta
Eosinophil
Myeloid dendritic cell activated
T cell CD4+ memory activated
Myeloid dendritic cell resting
Neutrophil
Mast cell resting
T cell CD4+ naive
Monocyte
Macrophage M2
Tumor type
CIBERSORT
0.2
0.15
0.1
0.05
0
ETMR
MNG
MB
SCHW
ATRT
EPN
pedHGG
pedLGG
NFB
CPH
CP
NBL

Figure S4

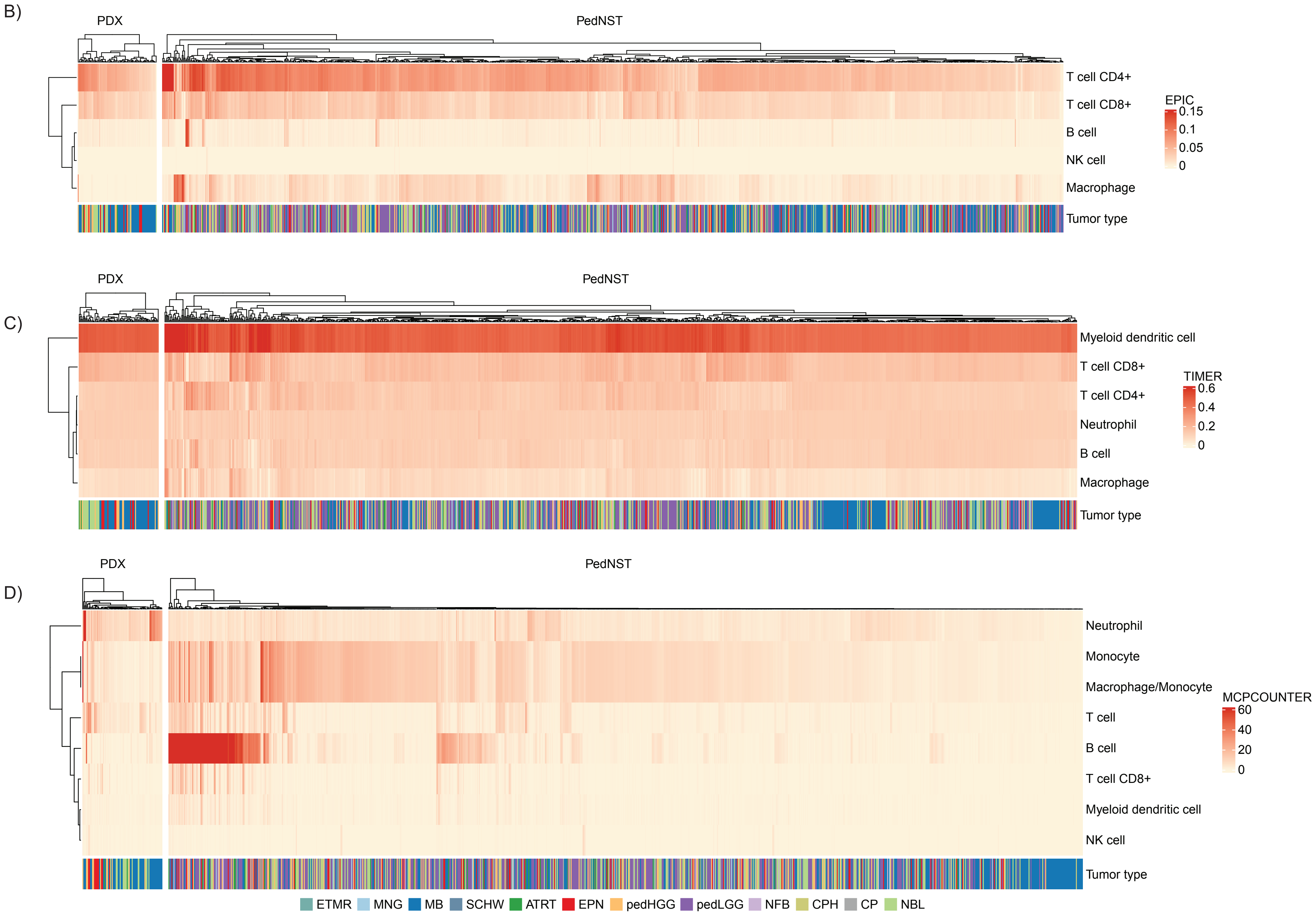
B)
PDX
PedNST
T cell CD4+
T cell CD8+
B cell
NK cell
Macrophage
Tumor type
EPIC
0.15
0.1
0.05
0
C)
PDX
PedNST
Myeloid dendritic cell
T cell CD8+
T cell CD4+
Neutrophil
B cell
Macrophage
Tumor type
TIMER
0.6
0.4
0.2
0
D)
PDX
PedNST
Neutrophil
Monocyte
Macrophage/Monocyte
T cell
B cell
T cell CD8+
Myeloid dendritic cell
NK cell
Tumor type
MCPCOUNTER
60
40
20
0
ETMR MNG MB SCHW ATRT EPN pedHGG pedLGG NFB CPH CP NBL

Figure S4

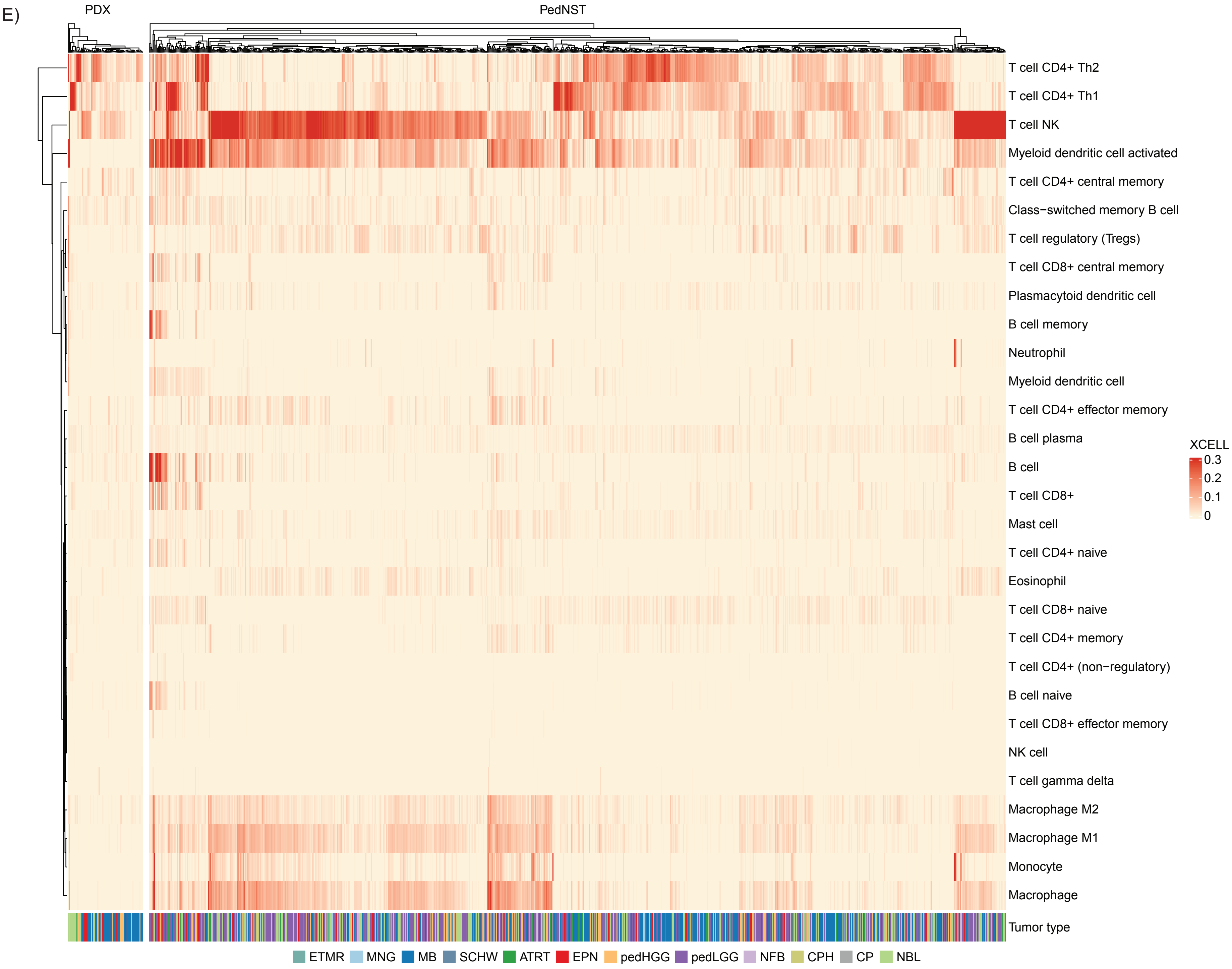
E)
PDX
PedNST
T cell CD4+ Th2
T cell CD4+ Th1
T cell NK
Myeloid dendritic cell activated
T cell CD4+ central memory
Class−switched memory B cell
T cell regulatory (Tregs)
T cell CD8+ central memory
Plasmacytoid dendritic cell
B cell memory
Neutrophil
Myeloid dendritic cell
T cell CD4+ effector memory
B cell plasma
B cell
T cell CD8+
Mast cell
T cell CD4+ naive
Eosinophil
T cell CD8+ naive
T cell CD4+ memory
T cell CD4+ (non−regulatory)
B cell naive
T cell CD8+ effector memory
NK cell
T cell gamma delta
Macrophage M2
Macrophage M1
Monocyte
Macrophage
Tumor type
XCELL
0.3
0.2
0.1
0
ETMR MNG MB SCHW ATRT EPN pedHGG pedLGG NFB CPH CP NBL

Figure S4

F)

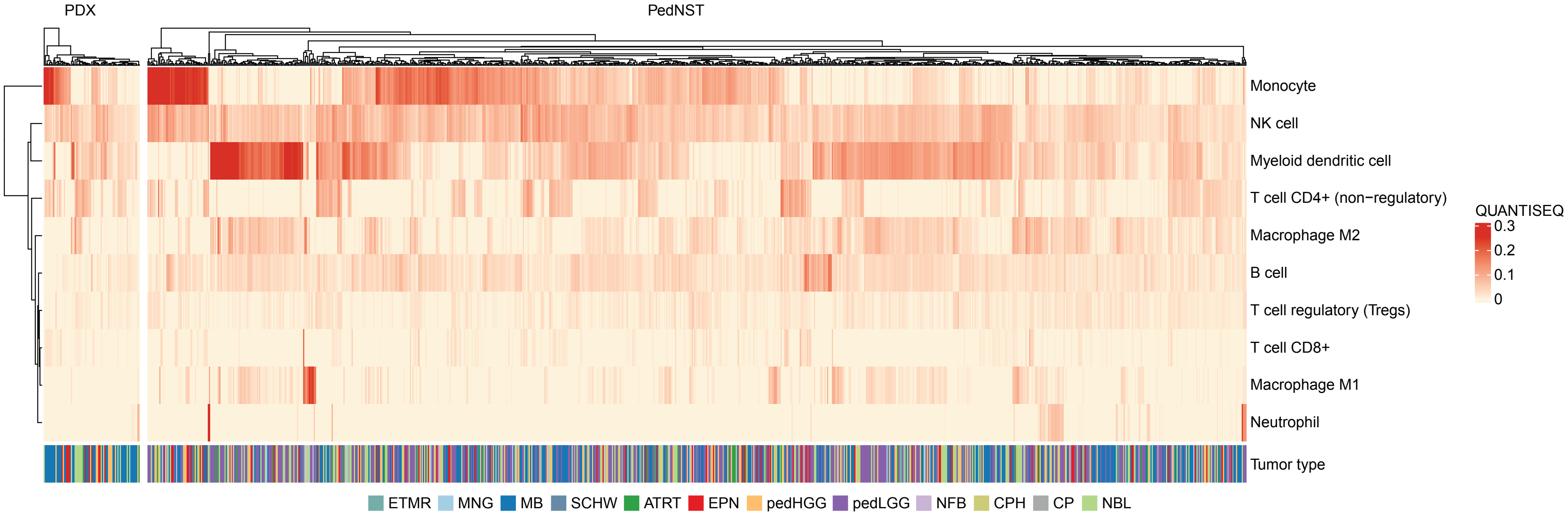
PDX
PedNST
Monocyte
NK cell
Myeloid dendritic cell
T cell CD4+ (non−regulatory)
Macrophage M2
B cell
T cell regulatory (Tregs)
T cell CD8+
Macrophage M1
Neutrophil
Tumor type
QUANTISEQ
0.3
0.2
0.1
0
ETMR
MNG
MB
SCHW
ATRT
EPN
pedHGG
pedLGG
NFB
CPH
CP
NBL

Figure S4

G)

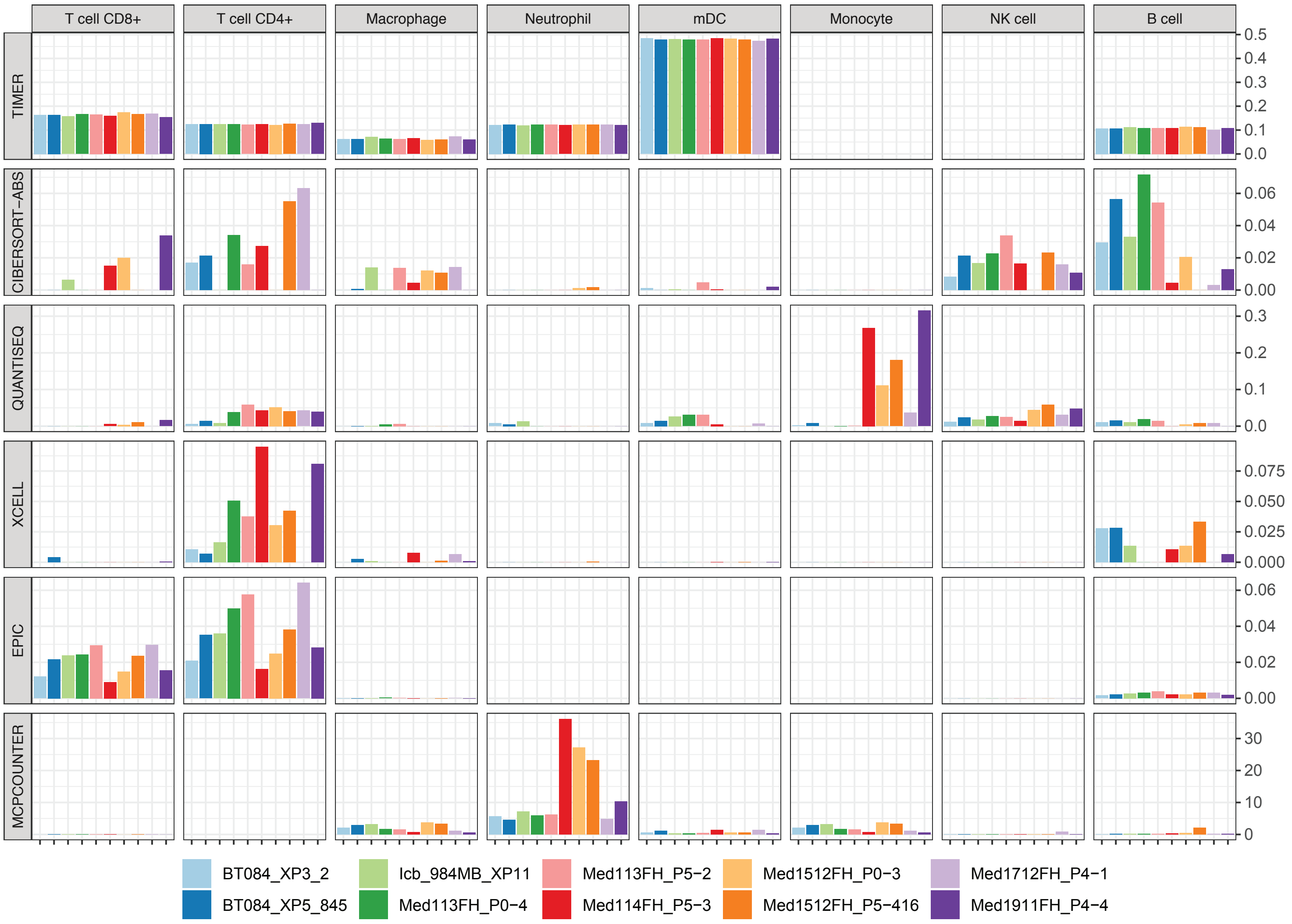

T cell CD8+
T cell CD4+
Macrophage
Neutrophil
mDC
Monocyte
NK cell
B cell
TIMER
CIBERSORT−ABS
QUANTISEQ
XCELL
EPIC
MCPCOUNTER
0.5
0.4
0.3
0.2
0.1
0.0
0.06
0.04
0.02
0.00
0.3
0.2
0.1
0.0
0.075
0.050
0.025
0.000
0.06
0.04
0.02
0.00
30
20
10
0
BT084_XP3_2
BT084_XP5_845
Icb_984MB_XP11
Med113FH_P0−4
Med113FH_P5−2
Med114FH_P5−3
Med1512FH_P0−3
Med1512FH_P5−416
Med1712FH_P4−1
Med1911FH_P4−4


Figure S4

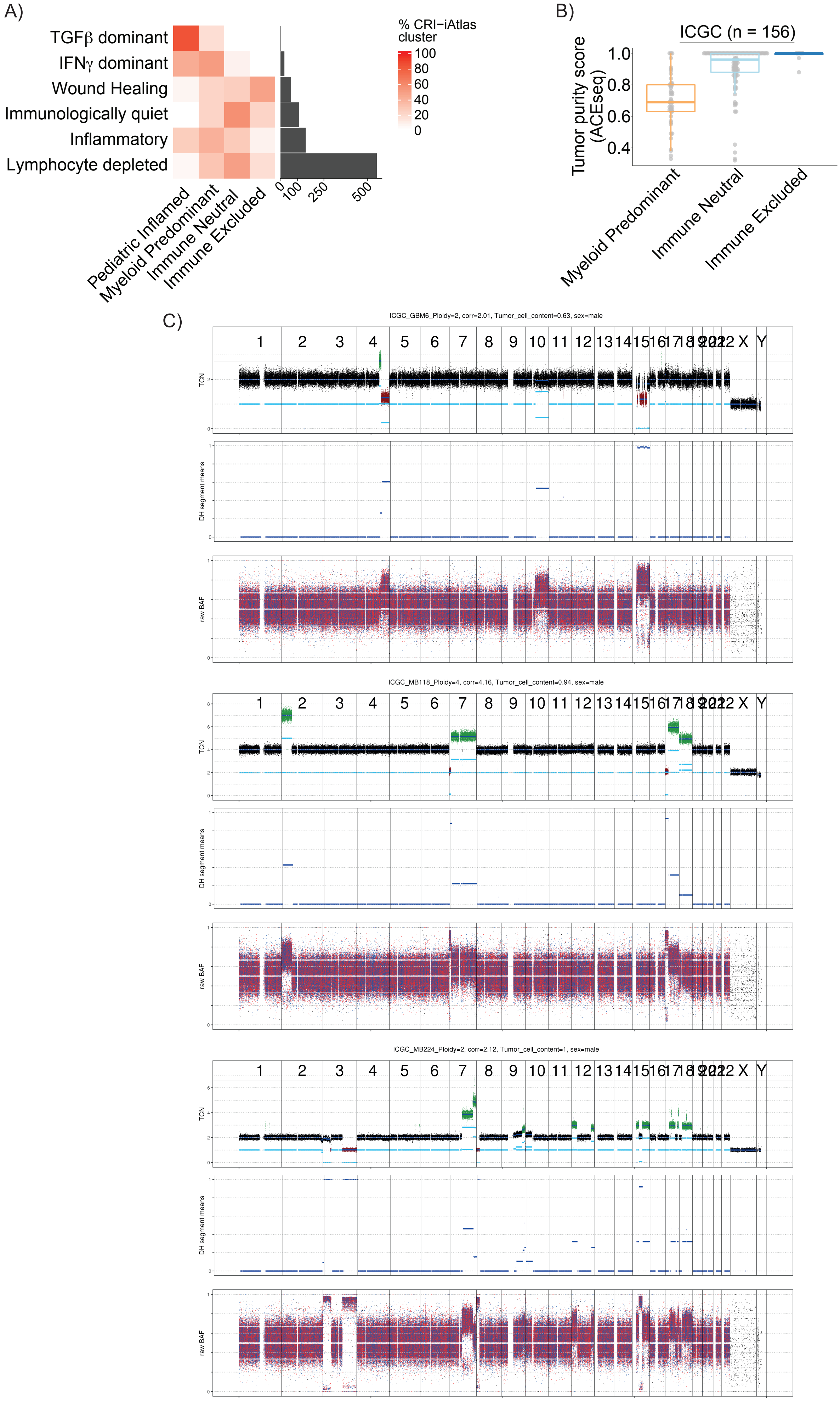
A)
TGFβ dominant
IFNγ dominant
Wound Healing
Immunologically quiet
Inflammatory
Lymphocyte depleted
Pediatric Inflamed
Myeloid Predominant
Immune Neutral
Immune Excluded
0
100
250
500
% CRI−iAtlas cluster
100
80
60
40
20
0
B)
ICGC (n = 156)
Tumor purity score (ACEseq)
1.0
0.8
0.6
0.4
Myeloid Predominant
Immune Neutral
Immune Excluded
C)
ICGC_GBM6_Ploidy=2, corr=2.01, Tumor_cell_content=0.63, sex=male
ICGC_MB118_Ploidy=4, corr=4.16, Tumor_cell_content=0.94, sex=male
ICGC_MB224_Ploidy=2, corr=2.12, Tumor_cell_content=1, sex=male
TCN
DH segment means
raw BAF

Figure S5

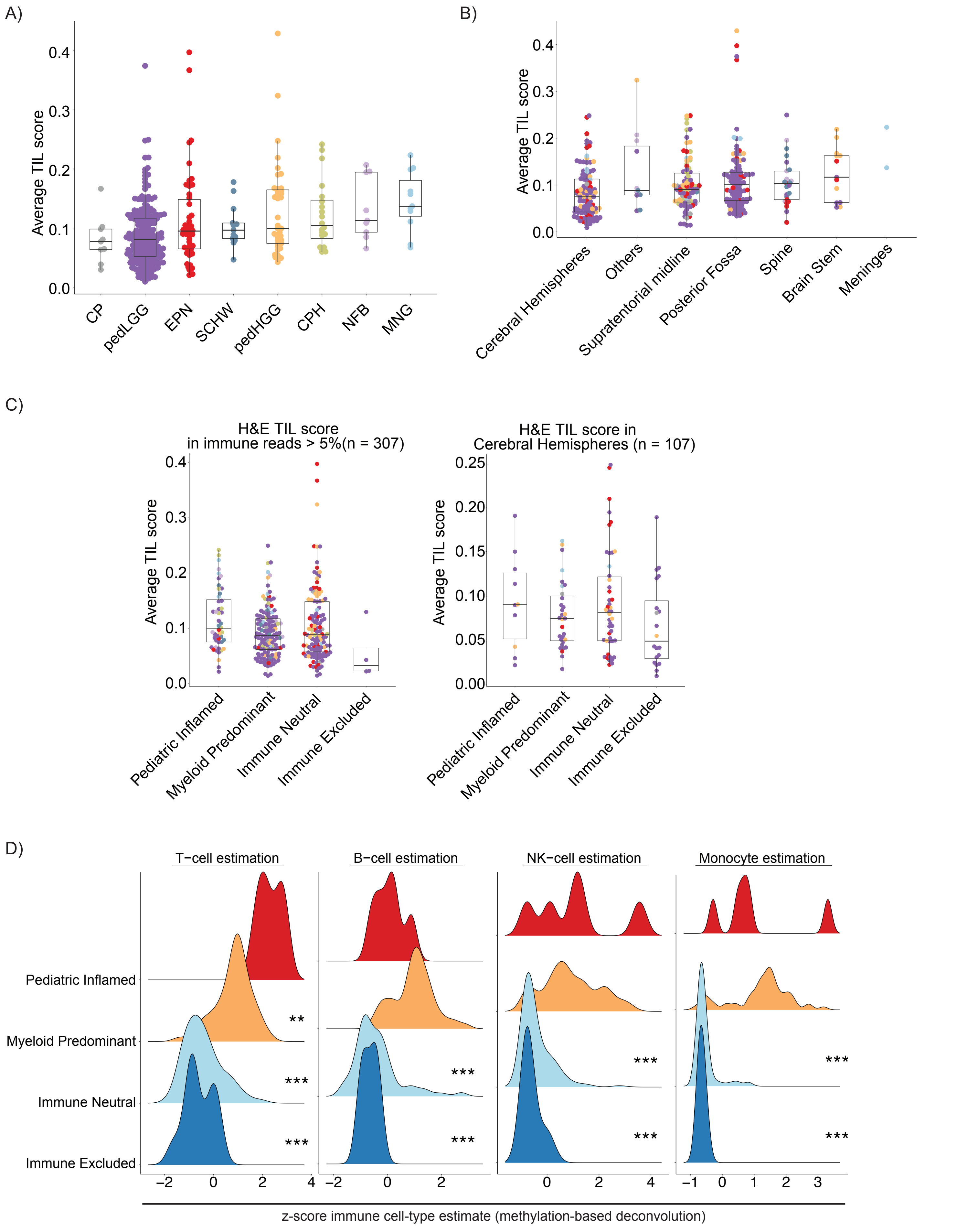
A)
Average TIL score
0.0
0.1
0.2
0.3
0.4
CP
pedLGG
EPN
SCHW
pedHGG
CPH
NFB
MNG
B)
Average TIL score
0.0
0.1
0.2
0.3
0.4
Cerebral Hemispheres
Others
Supratentorial midline
Posterior Fossa
Spine
Brain Stem
Meninges
C)
H&E TIL score
in immune reads > 5%(n = 307)
Average TIL score
Pediatric Inflamed
Myeloid Predominant
Immune Neutral
Immune Excluded
H&E TIL score in
Cerebral Hemispheres (n = 107)
Average TIL score
0.00
0.05
0.10
0.15
0.20
0.25
D)
T−cell estimation
B−cell estimation
NK−cell estimation
Monocyte estimation
Pediatric Inflamed
Myeloid Predominant
Immune Neutral
Immune Excluded
**
***
z-score immune cell-type estimate (methylation-based deconvolution)

Figure S6

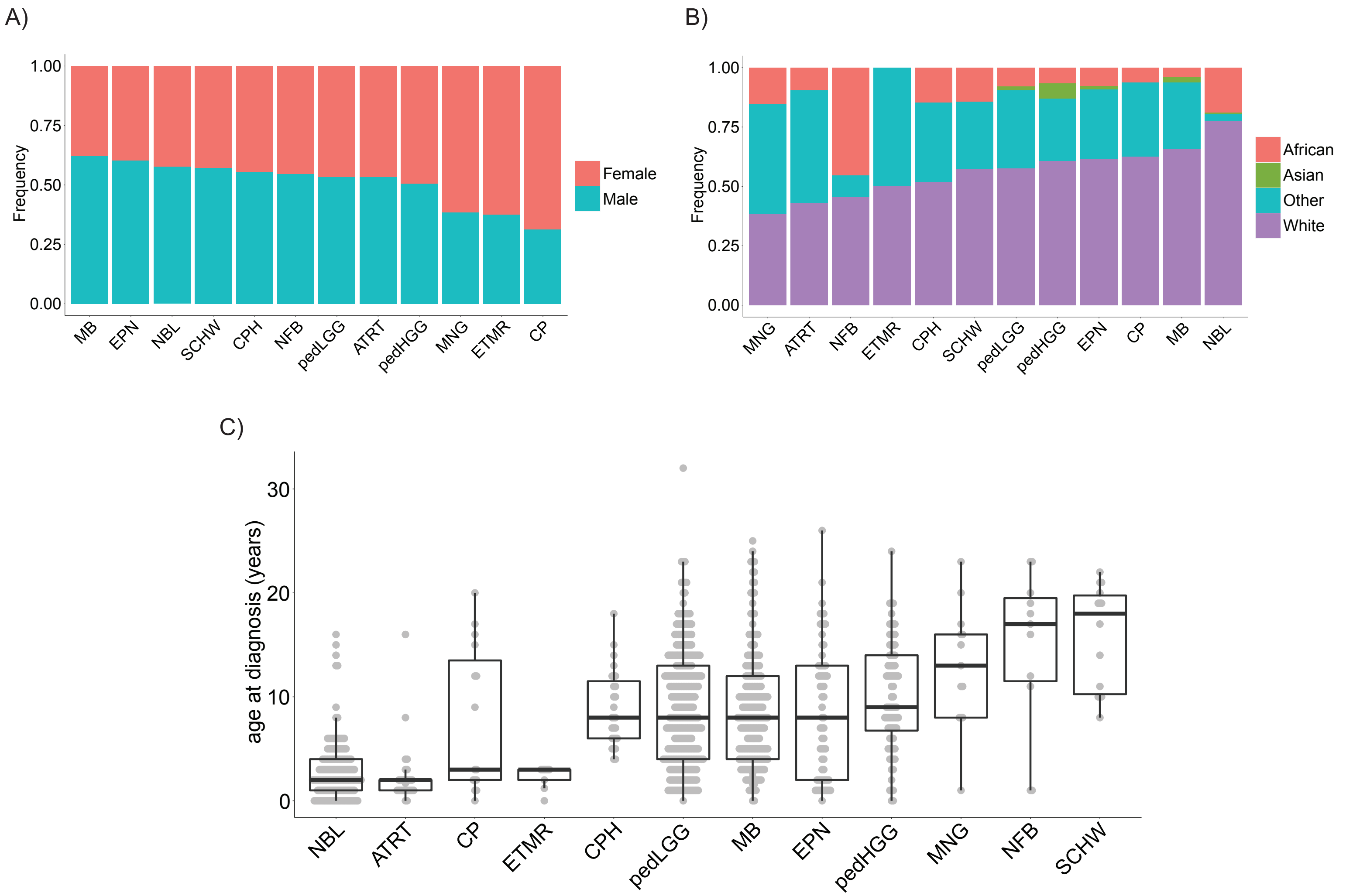

A)
Frequency
1.00
0.75
0.50
0.25
0.00
MB
EPN
NBL
SCHW
CPH
NFB
pedLGG
ATRT
pedHGG
MNG
ETMR
CP
Female
Male
B)
Frequency
1.00
0.75
0.50
0.25
0.00
MNG
ATRT
NFB
ETMR
CPH
SCHW
pedLGG
pedHGG
EPN
CP
MB
NBL
African
Asian
Other
White
C)
age at diagnosis (years)
30
20
10
0
NBL
ATRT
CP
ETMR
CPH
pedLGG
MB
EPN
pedHGG
MNG
NFB
SCHW


Figure S7

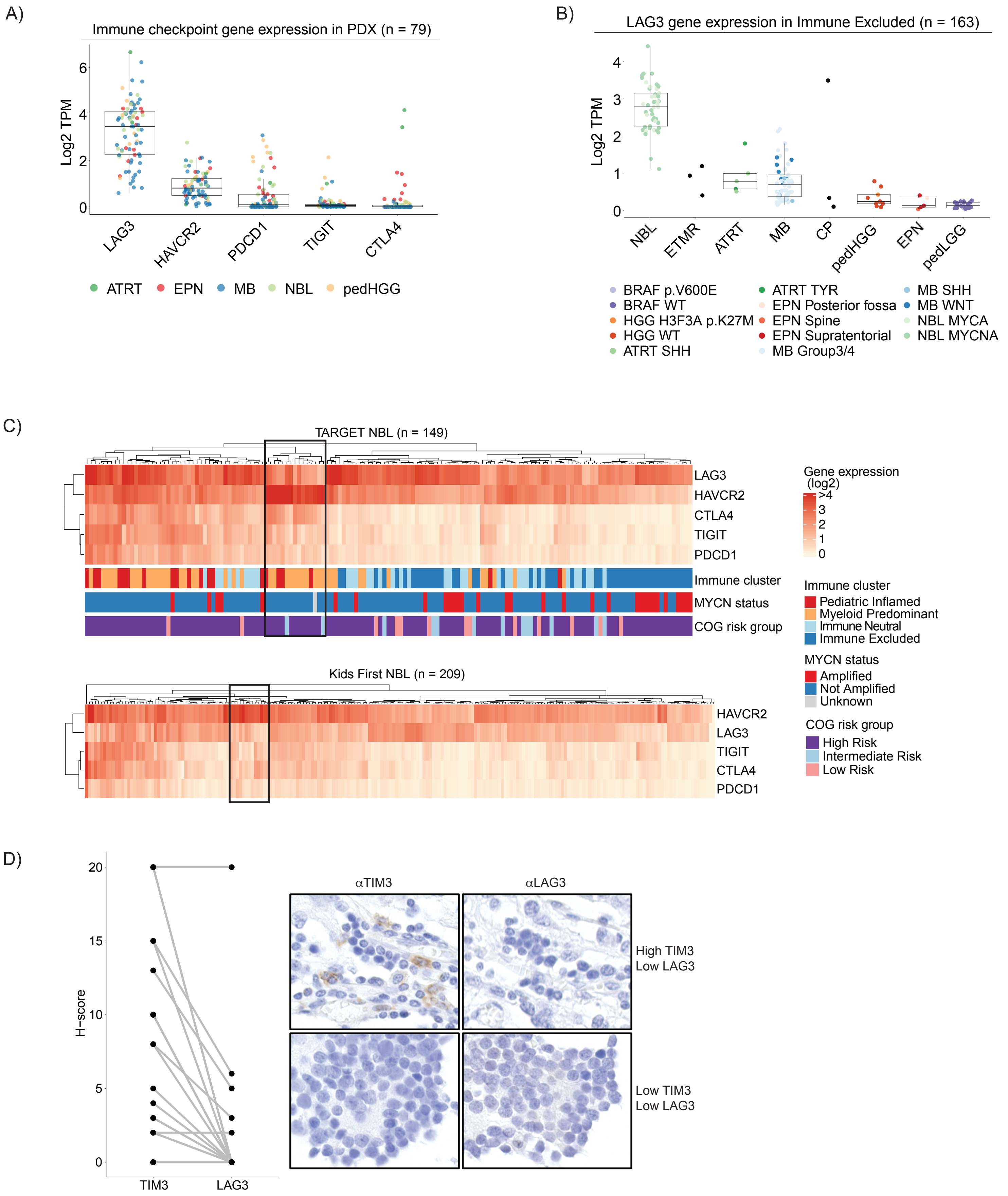
A)
Immune checkpoint gene expression in PDX (n = 79)
Log2 TPM
0
2
4
6
LAG3
HAVCR2
PDCD1
TIGIT
CTLA4
ATRT
EPN
MB
NBL
pedHGG
B)
LAG3 gene expression in Immune Excluded (n = 163)
Log2 TPM
0
1
2
3
4
NBL
ETMR
ATRT
MB
CP
pedHGG
EPN
pedLGG
BRAF p.V600E
BRAF WT
HGG H3F3A p.K27M
HGG WT
ATRT SHH
ATRT TYR
EPN Posterior fossa
EPN Spine
EPN Supratentorial
MB Group3/4
MB SHH
MB WNT
NBL MYCA
NBL MYCNA
C)
TARGET NBL (n = 149)
LAG3
HAVCR2
CTLA4
TIGIT
PDCD1
Immune cluster
MYCN status
COG risk group
Kids First NBL (n = 209)
HAVCR2
LAG3
TIGIT
CTLA4
PDCD1
Gene expression
(log2)
>4
3
2
1
0
Immune cluster
Pediatric Inflamed
Myeloid Predominant
Immune Neutral
Immune Excluded
MYCN status
Amplified
Not Amplified
Unknown
COG risk group
High Risk
Intermediate Risk
Low Risk
D)
H-score
0
5
10
15
20
TIM3
LAG3
αTIM3
αLAG3
High TIM3
Low LAG3
Low TIM3
Low LAG3

Figure S8

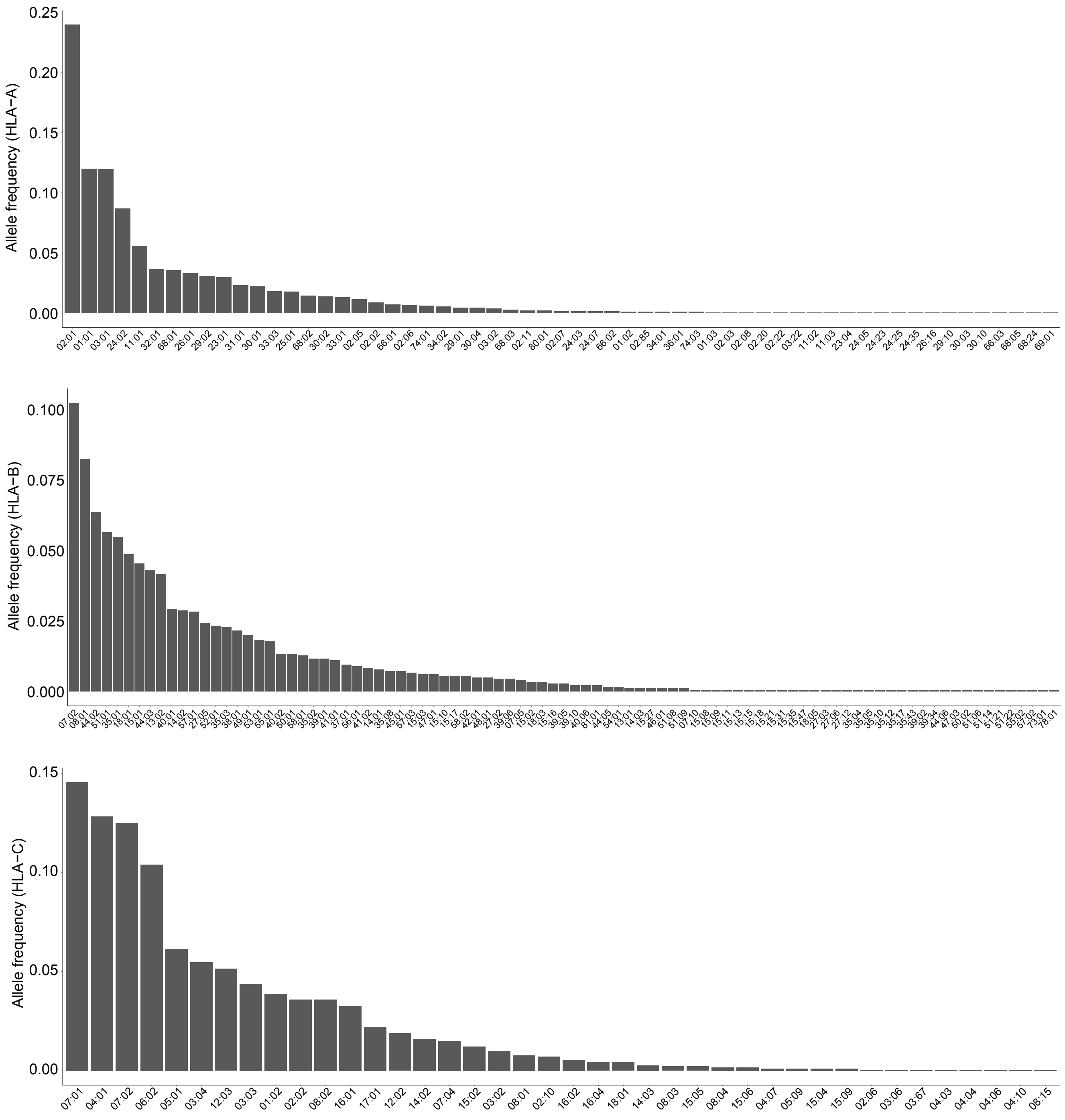

Allele frequency (HLA−A)
0.25
0.20
0.15
0.10
0.05
0.00
02:01 01:01 03:01 24:02 11:01 32:01 68:01 26:01 29:02 23:01 31:01 30:01 33:03 25:01 68:02 30:02 33:01 02:05 02:02 66:01 02:06 74:01 34:02 29:01 30:04 03:02 68:03 02:11 80:01 02:07 24:03 24:07 66:02 01:02 02:85 34:01 36:01 74:03 01:03 02:03 02:08 02:20 02:22 03:22 11:02 11:03 23:04 24:05 24:23 24:25 24:35 26:16 29:10 30:03 30:10 66:03 68:05 68:24 69:01
Allele frequency (HLA−B)
0.100
0.075
0.050
0.025
0.000
Allele frequency (HLA−C)
0.15
0.10
0.05
0.00
07:01 04:01 07:02 06:02 05:01 03:04 12:03 03:03 01:02 02:02 08:02 16:01 17:01 12:02 14:02 07:04 15:02 03:02 08:01 02:10 16:02 16:04 18:01 14:03 08:03 15:05 08:04 15:06 04:07 05:09 15:04 15:09 02:06 03:06 03:67 04:03 04:04 04:06 04:10 08:15


Figure S9

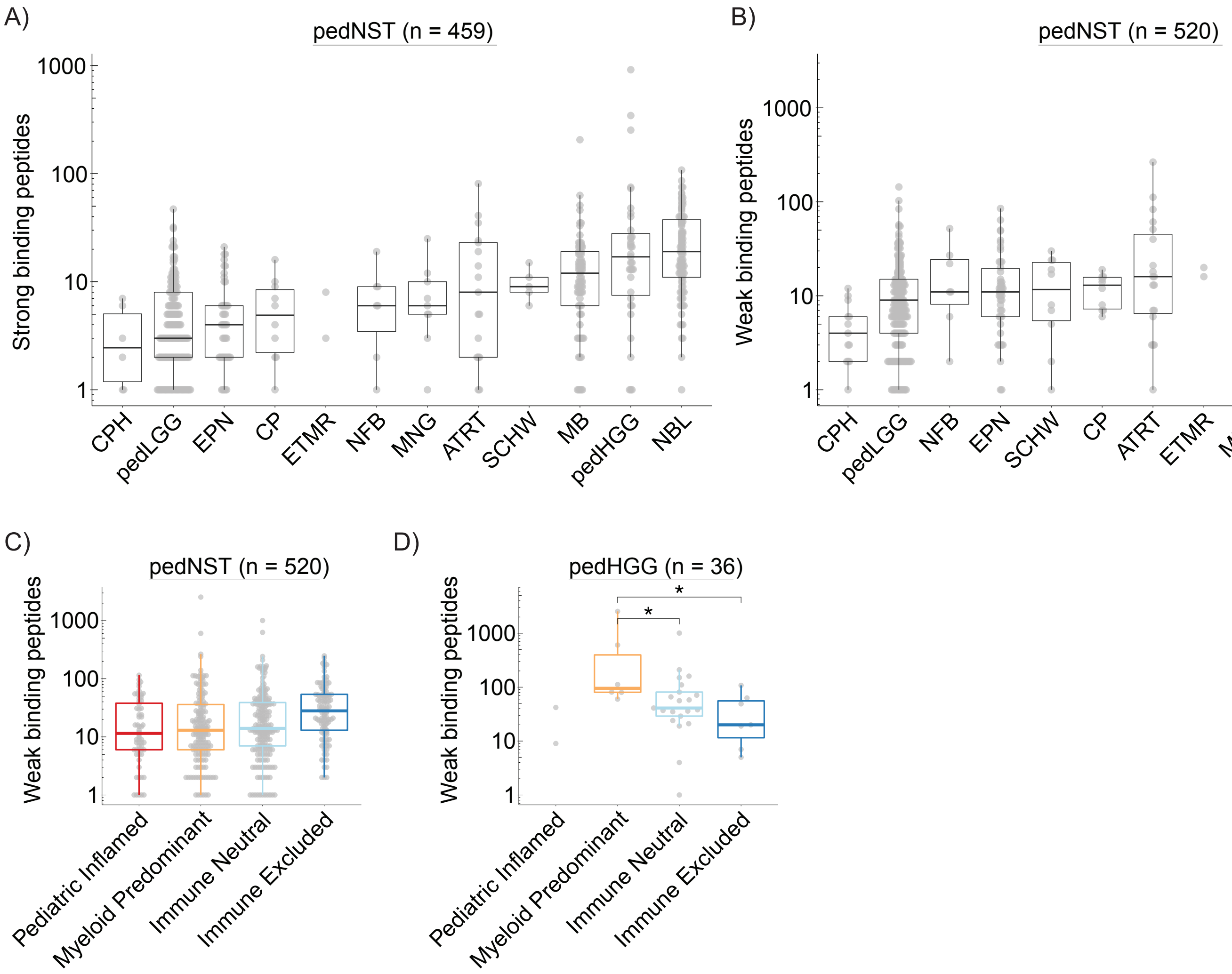
A)
pedNST (n = 459)
Strong binding peptides
1000
100
10
1
CPH
pedLGG
EPN
CP
ETMR
NFB
MNG
ATRT
SCHW
MB
pedHGG
NBL
B)
pedNST (n = 520)
Weak binding peptides
1000
100
10
1
CPH
pedLGG
NFB
EPN
SCHW
CP
ATRT
ETMR
MNG
MB
pedHGG
NBL
C)
pedNST (n = 520)
Weak binding peptides
1000
100
10
1
Pediatric Inflamed
Myeloid Predominant
Immune Neutral
Immune Excluded
D)
pedHGG (n = 36)
Weak binding peptides
1000
100
10
1
*
*
Pediatric Inflamed
Myeloid Predominant
Immune Neutral
Immune Excluded

Figure S10

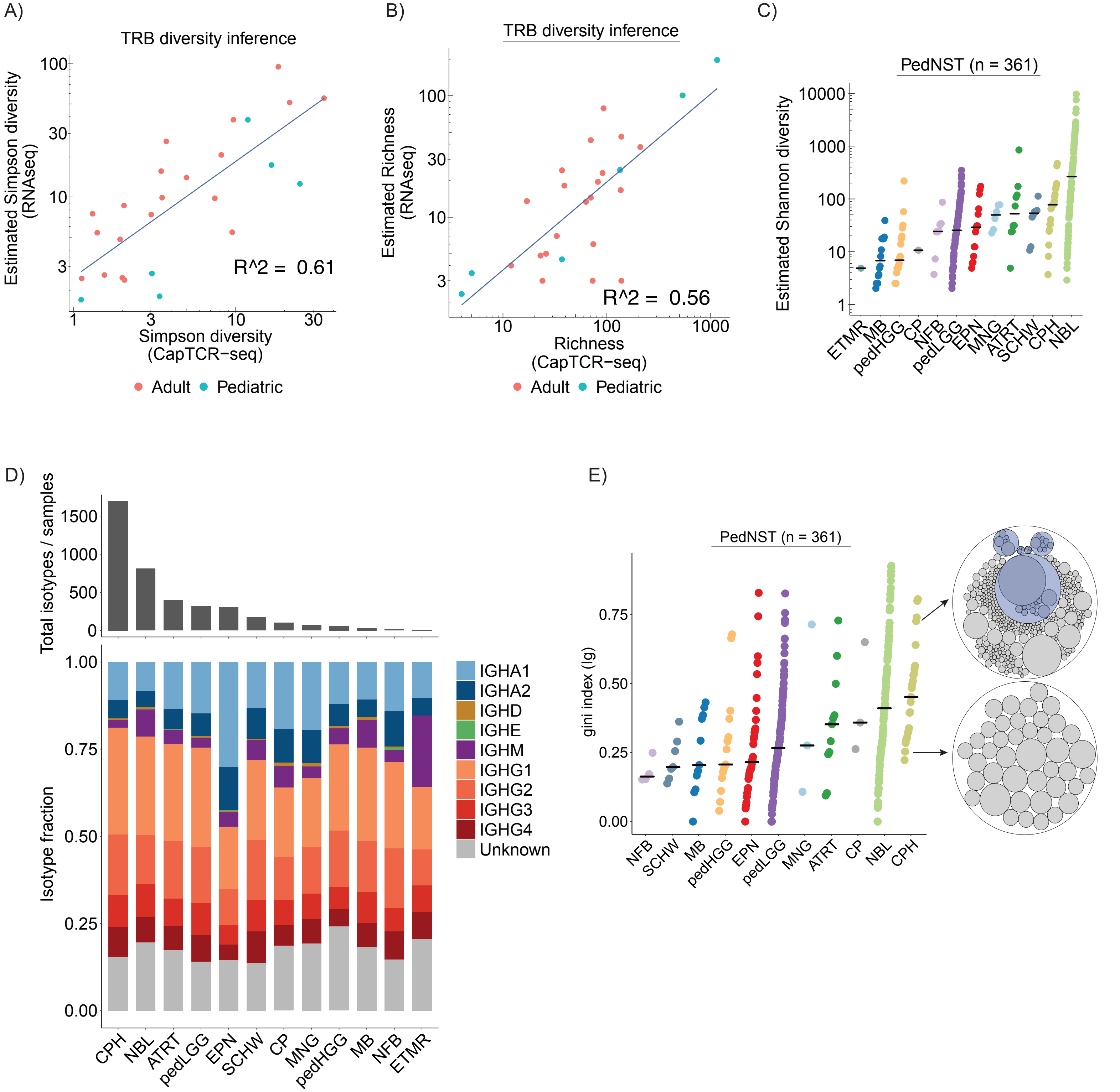
A)
TRB diversity inference
Estimated Simpson diversity (RNAseq)
Simpson diversity (CapTCR-seq)
R^2 = 0.61
Adult
Pediatric
B)
TRB diversity inference
Estimated Richness (RNAseq)
Richness (CapTCR-seq)
R^2 = 0.56
Adult
Pediatric
C)
PedNST (n = 361)
Estimated Shannon diversity
ETMR MB pedHGG CP NFB pedLGG EPN MNG ATRT SCHW CPH NBL
D)
Total isotypes / samples
Isotype fraction
IGHA1
IGHA2
IGHD
IGHE
IGHM
IGHG1
IGHG2
IGHG3
IGHG4
Unknown
CPH NBL ATRT pedLGG EPN SCHW CP MNG pedHGG MB NFB ETMR
E)
PedNST (n = 361)
gini index (lg)
NFB SCHW MB pedHGG EPN pedLGG MNG ATRT CP NBL CPH

Figure S11

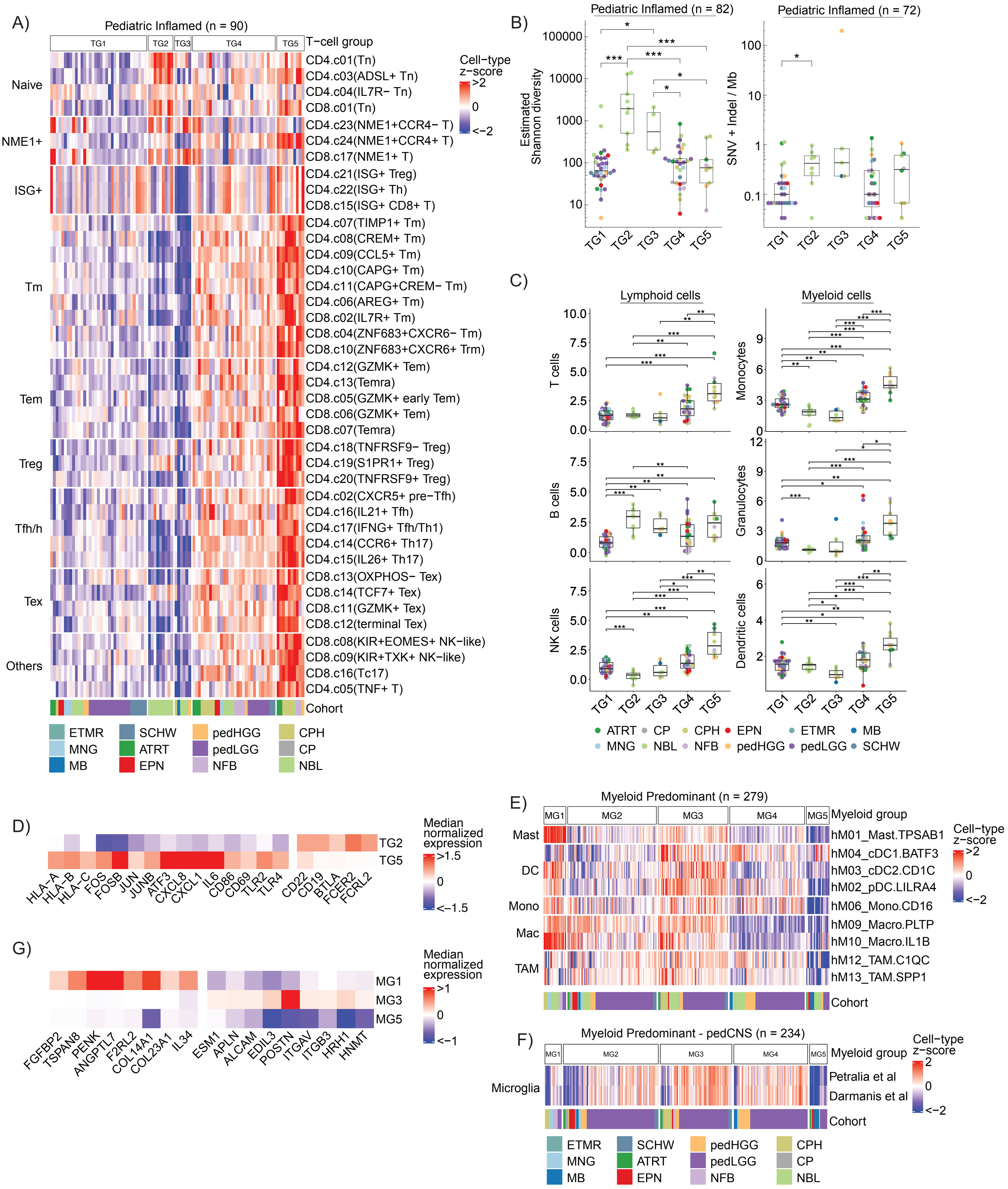
A)
Pediatric Inflamed (n = 90)
TG1 TG2 TG3 TG4 TG5 T−cell group
Naive
NME1+
ISG+
Tm
Tem
Treg
Tfh/h
Tex
Others
CD4.c01(Tn)
CD4.c03(ADSL+ Tn)
CD4.c04(IL7R− Tn)
CD8.c01(Tn)
CD4.c23(NME1+CCR4− T)
CD4.c24(NME1+CCR4+ T)
CD8.c17(NME1+ T)
CD4.c21(ISG+ Treg)
CD4.c22(ISG+ Th)
CD8.c15(ISG+ CD8+ T)
CD4.c07(TIMP1+ Tm)
CD4.c08(CREM+ Tm)
CD4.c09(CCL5+ Tm)
CD4.c10(CAPG+ Tm)
CD4.c11(CAPG+CREM− Tm)
CD4.c06(AREG+ Tm)
CD8.c02(IL7R+ Tm)
CD8.c04(ZNF683+CXCR6− Tm)
CD8.c10(ZNF683+CXCR6+ Trm)
CD4.c12(GZMK+ Tem)
CD4.c13(Temra)
CD8.c05(GZMK+ early Tem)
CD8.c06(GZMK+ Tem)
CD8.c07(Temra)
CD4.c18(TNFRSF9− Treg)
CD4.c19(S1PR1+ Treg)
CD4.c20(TNFRSF9+ Treg)
CD4.c02(CXCR5+ pre−Tfh)
CD4.c16(IL21+ Tfh)
CD4.c17(IFNG+ Tfh/Th1)
CD4.c14(CCR6+ Th17)
CD4.c15(IL26+ Th17)
CD8.c13(OXPHOS− Tex)
CD8.c14(TCF7+ Tex)
CD8.c11(GZMK+ Tex)
CD8.c12(terminal Tex)
CD8.c08(KIR+EOMES+ NK−like)
CD8.c09(KIR+TXK+ NK−like)
CD8.c16(Tc17)
CD4.c05(TNF+ T)
Cohort
Cell−type z−score
>2
0
<−2
ETMR SCHW pedHGG CPH
MNG ATRT pedLGG CP
MB EPN NFB NBL
B)
Pediatric Inflamed (n = 82)
Pediatric Inflamed (n = 72)
Estimated Shannon diversity
100000
10000
1000
100
10
SNV + Indel / Mb
100
10
1
0.1
TG1 TG2 TG3 TG4 TG5
C)
Lymphoid cells
Myeloid cells
T cells
B cells
NK cells
Monocytes
Granulocytes
Dendritic cells
10.0
7.5
5.0
2.5
0.0
12
9
6
3
0
TG1 TG2 TG3 TG4 TG5
ATRT CP CPH EPN ETMR MB
MNG NBL NFB pedHGG pedLGG SCHW
D)
TG2
TG5
HLA−A HLA−B HLA−C FOS FOSB JUN JUNB ATF3 CXCL8 CXCL1 IL6 CD86 CD69 TLR2 TLR4 CD22 CD19 BTLA FCER2 FCRL2
Median normalized expression
>1.5
0
<−1.5
E)
Myeloid Predominant (n = 279)
MG1 MG2 MG3 MG4 MG5 Myeloid group
Mast
DC
Mono
Mac
TAM
hM01_Mast.TPSAB1
hM04_cDC1.BATF3
hM03_cDC2.CD1C
hM02_pDC.LILRA4
hM06_Mono.CD16
hM09_Macro.PLTP
hM10_Macro.IL1B
hM12_TAM.C1QC
hM13_TAM.SPP1
Cohort
Cell−type z−score
>2
0
<−2
G)
MG1
MG3
MG5
FGFBP2 TSPAN8 PENK ANGPTL7 F2RL2 COL14A1 COL23A1 IL34 ESM1 APLN ALCAM EDIL3 POSTN ITGAV ITGB3 HRH1 HNMT
Median normalized expression
>1
0
<−1
F)
Myeloid Predominant - pedCNS (n = 234)
MG1 MG2 MG3 MG4 MG5 Myeloid group
Microglia
Petralia et al
Darmanis et al
Cohort
Cell−type z−score
2
0
<−2
ETMR SCHW pedHGG CPH
MNG ATRT pedLGG CP
MB EPN NFB NBL

Figure S12
